# Deep Variational Autoencoders for Population Genetics

**DOI:** 10.1101/2023.09.27.558320

**Authors:** Margarita Geleta, Daniel Mas Montserrat, Xavier Giro-i-Nieto, Alexander G. Ioannidis

## Abstract

**Motivation:** Modern biobanks provide numerous high-resolution genomic sequences of diverse populations. These datasets enable a better understanding of genotype-phenotype interactions with genome-wide association studies (GWAS) and power a new personalized precision medicine with polygenic risk scores (PRS). In order to account for diverse and admixed populations, new algorithmic tools are needed in order to properly capture the genetic composition of populations. Here we explore deep learning techniques, namely variational autoencoders (VAEs), to process genomic data from a population perspective. We hope this work will encourage the adoption of deep neural networks in the population genetics community.

**Results:** In this paper, we show the power of VAEs for a variety of tasks relating to the interpretation, classification, simulation, and compression of genomic data with several worldwide whole genome datasets from both humans and canids and evaluate the performance of the proposed applications with and without ancestry conditioning. The unsupervised setting of autoencoders allows for the detection and learning of granular population structure and inferring of informative latent factors. The learned latent spaces of VAEs are able to capture and represent differentiated Gaussian-like clusters of samples with similar genetic composition on a fine-scale from single nucleotide polymorphisms (SNPs), enabling applications in dimensionality reduction, data simulation, and imputation. These individual genotype sequences can then be decomposed into latent representations and reconstruction errors (residuals) which provide a sparse representation useful for lossless compression. We show that different population groups have differentiated compression ratios and classification accuracies. Additionally, we analyze the entropy of the SNP data, its effect on compression across populations, its relation to historical migrations, and we show how to introduce autoencoders into existing compression pipelines.

## 1 Introduction

Deep learning is becoming ubiquitous across all areas of science and engineering, with artificial neural networks (ANNs) being used to model highly non-linear and complex data and proving successful in a wide range of applications. Recent works have begun to introduce such techniques within the fields of population genetics and precision medicine [1–3]. Here we explore the use of variational autoencoders (VAEs), a type of neural network used to learn a low-dimensional representation of the data, to analyze sequences of single nucleotide polymorphisms (SNPs), and showcase many applications including dimensionality reduction, data compression, classification, and simulation.

The number of human genomes being sequenced every year is growing rapidly, fueled by dramatic improvements in sequencing technology. Biobanks, paramount in areas like precision medicine, are powering genome-wide association studies (GWAS), where many genetic variants (e.g. SNPs) are analyzed across different subjects to find the relationships between genetic and phenotypic traits [4, 5], and are used to develop new treatments and drugs [6]. This creates a need for new, efficient and accurate data-driven algorithmic tools to store, visualize and characterize high-dimensional genomic data. While traditional statistical techniques used to analyze genetic data become computationally expensive and slow when faced with biobank-sized data, neural networks can provide a powerful alternative.

### Ancestry Prediction

The relationship between ancestry, ethnicity, race, and genetic variation remains controversial, involving biology, ethics, and biopolitics. Because genetic variation might not follow the racial identities established culturally and historically [7], data-driven alternatives are needed and remain an active area of research. The term *ancestry* refers to a category of genetic similarity that can be associated with a shared origin, social background, cultural heritage, and environment [8, 9]. The framework for classifying or regressing genetic ancestry translates these ancestry effects from macro-level biological differences to the molecular level of genes [10]. Indeed, an individual’s geographic origin can be inferred with remarkable accuracy from their DNA [11]. This task, known as *global ancestry inference*, can be either approached from a discrete perspective (classifying the ancestry label) or from a continuous one (regressing the geographical coordinates of origin). Some widely adopted techniques include ADMIXTURE [12, 13], a clustering technique based on probabilistic non-negative matrix factorization, and its more recent neural counterpart – Neural ADMIXTURE [3]; Locator [14], a *multi-layer perceptron* (MLP) that addresses the *geographical coordinate regression* problem by estimating a nonlinear function mapping genotypes to locations, and Diet Networks [2], which represent another deep-learning-based ancestry classifier within this paradigm of methods. Nevertheless, characterizing ancestry as a set of discrete, predefined labels can be limiting, as an increasing number of individuals are admixed, stemming from ancestors belonging to multiple ancestral population groups. For instance, the genomes of numerous African-Americans, have variable proportions of segments that could be classified as of European and West African ancestry [15]. State-of-the-art *local ancestry inference* (LAI) methods, rely on neural networks as well – LAI-Net [1] and SALAI-Net [16] replicate the classification-smoothing paradigm of RFMix [17] but from a neural-based viewpoint.

### Genomic Sequences Compression

The storage and transmission of high-dimensional biobank data require substantial storage space and channel capacity. This need has motivated the development of high-performance compression tools tailored to the unique particularities of this type of data [18–20], which include the inherently high dimensionality of the data and its sparse yet complex structure. The demand for more powerful compression approaches becomes increasingly vital in modern large biobanks that contain genomic sequences produced by next-generation sequencing.

ANN-based approaches had already shown superior compression ratios decades ago [21, 22]. The first neural compressors mimicked the adaptive *prediction by partial matching* (PPM) model, where a neural network is used to estimate (and/or adjust) the probabilities for each symbol and then a usual coding method such as arithmetic coding is used to convert the data into a compressed bit-stream [22]. Following this line, *DeepZip* [23] estimated the probabilities by processing genomic sequences with GRUs and LSTMs. Mahoney introduced *logistic mixing* [24], a neural network without hidden layers that uses a simple update rule to adjust the output probabilities for better bit-stream coding. This idea has been brought to genomic compression of sequencing reads with *GeCo3* [25]. Another recent use of neural networks for DNA compression has been shown in *DeepDNA* [26], trained specifically on mitochondrial DNA data, using a convolutional layer to capture local features which are then combined and fed into a recurrent layer to output the probabilities for each symbol. On the other hand, autoencoders present an alternative to PPM. To the authors’ knowledge the only attempt to compress genomic data with autoencoders has been performed on sequencing reads data [27].

### Genotype Imputation

Missing data at several genomic positions can arise due to sequencing errors. Furthermore, many applications prefer low-resolution genotype arrays over high-resolution whole genome sequencing technologies due to cost considerations. This situation underscores the need for accurate and efficient imputation tools capable of predicting missing SNP positions [28]. Genotype imputation infers missing values by using correlations between SNPs induced by *linkage disequilibrium* (LD) [29]. Such imputation algorithms have typically been based on *hidden Markov models* (HMMs), which capture the LD in the sequences and try to infer the missing values. An example of such a model is *Beagle* [30]. More recently, ANNs have been introduced into the genomic data imputation field, with the use of fully-connected variational [31] and convolutional [32] denoising autoencoder models.

### Genomic Sequences Simulation

While the number of sequenced genomes has grown substantially over the years, there is a clear disparity among the ancestries represented. The proportion of participants of non-European descent has remained constant [4], potentially introducing bias toward European genomes and giving rise to the “missing diversity” problem [33]. To illustrate, as of 2018, the majority of GWAS encompassed approximately 78% individuals of European ancestry. Additionally, certain communities, predominantly comprised of individuals with non-European ancestry, are reluctant to participate in genetic studies due to privacy concerns or apprehensions about potential misuse, as seen in prior cases [34, 35]. To circumvent these challenges, data simulation tools can be used to augment genomic databases and offer mechanisms for sharing synthetic data possessing equivalent statistical properties, all while safeguarding the privacy of individuals. Several recent studies have explored the effectiveness of generative deep neural networks in generating simulated genotypes. Montserrat et al. use a class-conditional VAE-GAN to generate artificial yet realistic genotypes [36], while Yelmen et al. generate high-quality synthetic genomes with GAN and RBM [37]. Moment matching networks have provided competitive results for data simulation [38]. Furthermore, another work [39] has attempted to use VAEs for genotype simulation.

## Methods

### Human dataset

In this work, we employ a dataset comprising multiple sequences derived from publicly available human whole genome sequences collected from diverse worldwide populations. The three sources are: (1) *The 1000 genomes project* [40], reporting genomes of 2,504 individuals from 26 populations from all continents; (2) *The Human Genome Diversity Project* [41], adding 929 diverse genomes from 54 geographically, linguistically, and culturally diverse human populations; (3) *The Simons Genome Diversity Project* [42], providing genomes from 300 individuals from 142 diverse populations. The dataset is pruned to contain only single-ancestry origin individuals, i.e., individuals whose four grandparents self-reported belonging to the same ancestral group. After pruning, the dataset has 2,965 single-ancestry phased human genomes, each containing a maternal and paternal copy, resulting in a total of 5,930 haploid sequences, referred to as *founders*. Founders are split into three non-overlapping groups with proportions 80%, 10% and 10%, to generate the training, validation, and test sets, respectively. Following the simulation scheme outlined in [1], for each dataset, we simulate samples with the corresponding set of founders via online *Wright-Fisher* simulation [17, 43], where, at each VAE forward step, the online simulator produces new samples on-the-fly for each population separately, basing the recombination on the human *HapMap* genetic map [44]. To construct whole genome sequences, we apply the following steps independently to each chromosome: (1) *minor allele frequency* (MAF) filtering, i.e., we remove SNP positions with a MAF inferior than 0.01, and (2) LD pruning, i.e., we eliminate each SNP that has a *R*^2^ pearson coefficient value larger than 0.01 with any other SNP in a 50-SNP sliding window with a 10-SNP hop size. In each simulated batch, we ensure an equal number of individuals are generated for each population group, preventing training bias toward any specific ancestry. For the task of dimensionality reduction, we simulate samples within each of the 55 human sub-populations disjointly, so that there are no admixtures among subpopulations. Refer to the **Appendix** 1 for the full subpopulations list.

### Canine dataset

The canine dataset consists of 722 canine whole genome sequences sourced from [45]. For this dataset, we sample the most variable SNP positions among breeds across 38 canine chromosomes, which correspond to a subset of SNPs that matches the genotyping array used by the commercial brand Embark [46]. This dataset encompasses a diverse range of canids, documenting wild canids, indigenous and village dog populations, as well as 144 domestic dog breeds. To manage the extensive breed diversity, we group similar breeds into 15 distinct clades, as outlined in **Appendix** 1.

### Variational Autoencoders

Representation learning, also known as *feature learning*, attempts to recover a compact set of so-called latent *q*-dimensional variables ***z*** that describe a distribution over the *d*-dimensional observed data ***x***, with *q < d. Principal component analysis* (PCA) is a well-established statistical procedure for dimensionality reduction. For a set of observed data ***x***, the latent variables ***z*** are the orthonormal axes onto which the retained variance under projection of the data points is maximal. PCA can be given a natural probabilistic interpretation as the dimensionality reduction process can be considered in terms of the distribution of the latent variables, conditioned on the observation [47], where, from factor analysis, the relationship between ***x*** and ***z*** is linear and, conventionally, Gaussianity assumptions are taken. Autoencoders can be seen as a nonlinear generalization of PCA [48]. There are cases where the relationship between ***x*** and ***z*** is not linear and the common simplifying assumptions about the Gaussianity of the marginal or posterior probabilities do not reflect real data. In those cases, autoencoders are a perfect fit since they learn a direct encoding – a parametric map from inputs ***x*** to their representation latent ***z***. In this setup, two closed-form parametrized functions are defined: (1) the *encoder* 𝒱_*e*_ : R^*d*^ → R^*q*^, and (2) the *decoder* 𝒱_*d*_ : R^*q*^ → R^*d*^. Both 𝒱_*e*_(·) and 𝒱_*d*_(·) can be as expressive as desired: from a single linear layer to a MLP, or any other ANN architecture. In VAEs, the input ***x*** ∈ ℝ^*d*^ is encoded with 𝒱_*e*_(·) into the mean ***μ*** and a function of the variance ***σ*** vectors. Reparametrizing, the latent representation ***z*** = ***μ*** + ***σ*** ⊙ ***ϵ*** is obtained, with ***ϵ*** ∼ 𝒩 (**0, *I***), ⊙ being the element-wise product, and ***z, μ, σ, ϵ*** ∈ ℝ^*q*^. The latent representation can then be decoded with the decoder 𝒱_*d*_(·) to obtain the reconstruction 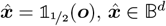, where ***o*** = 𝒱_*d*_(𝒱_*e*_(***x***)) ∈ ℝ^*d*^ is the output of the network and 1_1*/*2_ (·) is a unit step function applied elementwise on the output, binarizing each element of the output with a threshold of 1*/*2. We define the composition of the encoder-decoder pair and the binarization layer as 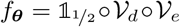, with parameters ***θ***, such that 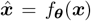, and since our input is always a sequence of binary values we have *f*_***θ***_ : 𝔹^*d*^ → 𝔹^*d*^. As the latent representation, located at the *bottleneck* of the architecture between the encoder and decoder, has a smaller dimensionality than the input (*q < d*), the primary learning objective is to compel the encoder to preserve as much of the relevant information as possible within this limited dimensionality.

Autoencoders learn the mapping function from input to feature space (the space spanned by ***z***) and the reverse mapping without learning a probability distribution of the data. In contrast, VAEs learn a probability distribution of the data [49] by enforcing an isotropic Gaussian prior over the latent variables. Since they learn to model the data, new samples can be generated by sampling, meaning that VAE are *generative* autoencoders.

### VAE architecture

Given the high dimensionality of genomic data and the data redudancy present from LD, autoencoders are exceptionally well-suited for this context. They excel in the task of learning novel, meaningful, and compact representations of SNP sequences, which represent the most prevalent genetic variations. The proposed VAE consists of a highly-adaptable and modular architecture, designed to accommodate different modes by utilizing flags for conditioning and denoising. Additionally, the model accepts two sets of parameters: (1) a set of *fixed parameters*, which defines essential aspects such as the number and size of layers in the encoder/decoder, dropouts, batch normalization and activation functions, and (2) a set of *hyperparameters*, which defines optimizer-related flags and values, including the learning rate *α*, variational *β*, weight decay *γ*, among others. The proposed VAE is composed of two symmetric MLP sub-networks: the encoder and the decoder. Both blocks consist of a stack of fully-connected layers. In the encoder, the layers progressively reduce in size, meaning each subsequent layer contains fewer neurons. Conversely, in the decoder, the layers expand, with each layer having a greater number of neurons in sequence. At the bottleneck of the architecture we find two feature maps: one trained to be the mean vector and another one the log-variance vector. These two feature maps, in conjunction with the VAE reparametrization trick, combine to yield the latent representation of the input ***z***. At each layer within the encoder, we enforce all units to conform to a Gaussian distribution by applying batch normalization [50]. This normalization operation serves two primary purposes: first, it encourages the network to shape the latent variable ***z*** as a Gaussian distribution, and second, it enhances the flow of gradients throughout the network. Following each fully connected layer, a non-linearity is applied. We have experimented with a full set of different non-linearities, and the best results are achieved with *rectified linear units* (ReLUs) [51] and *Gaussian error linear units* (GELUs) [52], in that order. The final layer of the decoder utilizes the sigmoid activation function to produce the probability vector for SNP positions, denoted as ***o***. These resulting probabilities are clamped with a unit step function 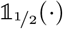 applied elementwise to obtain the reconstruction of the input 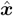. While our implementation allows the adjustment of the shape of the network, our current default settings use three dense layers of 512, 64 and 2 neurons, respectively, for the task of dimensionality reduction, and two dense layers of 512 and 64 neurons for all of the other tasks.

### Loss function

The loss function employed for training the VAE encompasses two primary objectives: the generative loss term and the latent loss term.

The generative loss objective strives for a high-quality reconstruction, aiming to make 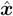 as close as possible to ***x***, so that 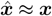. When we seek to maximize the probability of a reconstructed SNP position *x*_*i*_, 1 ≤ *i* ≤ *d* belonging to a known distribution with specific distribution parameters, our goal is to maximize the likelihood function for that particular distribution. In the **Results** section, we explain that each individual SNP can be effectively modeled with a Bernoulli distribution. Therefore, maximizing the likelihood for a Bernoulli distribution translates to minimizing the cross-entropy loss between ***x*** and ***o***.

Let ***θ*** represent the parameters of the VAE model, denoted as *f*_***θ***_(·), and let us denote the dataset 𝒟_*x*_ = {***x***_*n*_ | 1 ≤ *n* ≤ *N }*as the set of *d*-dimensional samples. Assuming i.i.d. data samples from the distribution *p*(𝒟_*x*_), we can define a multinomial likelihood in the following form:

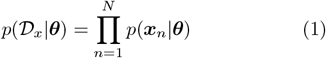

Under the assumption that each dimension of the SNP array is independent, we can approximate the likelihood function as follows:

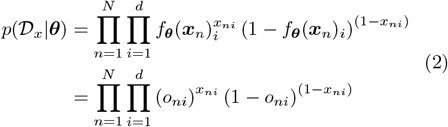

Taking the logarithm on this expression, we obtain:

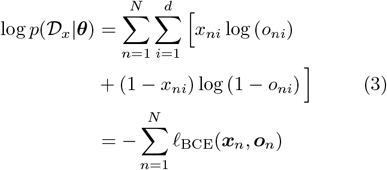

Thus, the generative loss for the VAE is computed as the *binary cross-entropy* (BCE) loss, *𝓁*_*BCE*_, between each sample ***x***_*n*_ and the VAE output ***o***_*n*_ for that sample.

The latent loss enforces that ***z*** follows a standard Gaussian distribution, i.e., ***z∼𝒩***(**0, 1**). To compare the distribution of the latent vector with a zero-mean, unit-variance Gaussian distribution, we employ the *Kullback-Leibler* (KL) divergence. This results in 𝒟_*KL*_ (p(z|x)||𝒩 (0, 1)) where z|x ∼ 𝒩(μ,Σ) is the encoder distribution defined by the two vectors at the bottleneck: the mean vector ***μ*** and the logarithm of the variance vector diag(**Σ**), as **Σ** is assumed to be a diagonal covariance matrix. The latent loss acts as a regularizing term, which is weighted with variational *β*. The loss for a given input ***x*** and VAE-computed output ***o*** is given by:

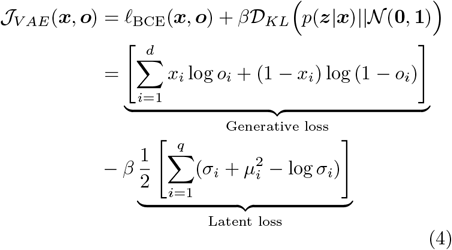

where *d* denotes the dimensionality of the observed data, ***x***, while *q* represents the dimensionality of the latent representation, ***z***. In our experiments, the land-scape of ℒ_*V AE*_ is traversed by the quasi-hyperbolic momentum variant of Adam (QHAdam) optimizer [53].

### Ancestry-conditional VAEs

Note that the above approach does not condition VAE on ancestry labels during training – it trains on all populations together. However, an ancestry-conditioned VAE includes the additional information of the population label *y*_*n*_ for each input sample ***x***_*n*_. This means we need to account for the labeled dataset 𝒟 = (𝒟_*x*_, 𝒟_*y*_) in the loss function, where dataset 𝒟_*x*_ = {***x***_*n*_|1 ≤ *n* ≤ *N* } is the set of *d*-dimensional samples and 𝒟_*y*_ = {*y*_*n*_|1 ≤ *n* ≤ *N, y*_*n*_ ∈ 𝒴} is the set of corresponding ancestry labels, where is 𝒴 the set of populations present in the data. After incorporating ancestry conditioning in Equation 1, and following a similar derivation as in previous Equations 2 and 3, we arrive at the same generative loss objective for the ancestry-conditioned VAE:

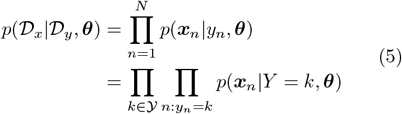

This indicates that the objective for the generative loss remains consistent in the ancestry-conditioned VAE:

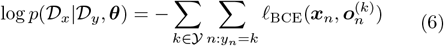

denoting the VAE output conditioned on ancestry label *k* as 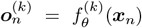. In practical terms, there are different ways to incorporate conditioning on the *k*-_*ed*_ th ancestry into both the encoder and the decoder, which we denote as 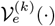 and 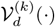, respectively. One approach, which we refer to as *regular* C-VAE (conditioned VAE), appends a one-hot encoded ancestry label to of both the encoder and the decoder. An alternative approach for conditioning fits a separate VAE for each population group individually, resulting in ancestry-specific overfitted VAEs. We refer to this approach as Y-VAE (Y-overfitted VAE), a method which has |Y| number of times more parameters than a regular VAE.

### Bayesian-motivated classification objectives

When employing ancestry conditioning on a VAE, an ancestry label can be inferred through *maximum a posteriori* (MAP) estimation. The output of the VAE conditioned on ancestry *k* is a vector containing Bernoulli probabilities 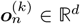. To infer an ancestry label, this vector is thresholded with a value greater than ½, resulting in 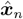. Because each SNP position is independently modeled as a Bernoulli distribution, the joint distribution can be expressed as the product of individual SNP distributions. Applying Bayes’ rule, we can derive *p*(*Y* = *k*|***x***_*n*_) ∝ *p*(***x***_*n*_|*Y* = *k*)*p*(*Y* = *k*), where *p*(***x***_*n*_ *Y* = *k*) represents the Bernoulli likelihood and the prior *p*(*Y* = *k*), for simplicity, is defined as the categorical distribution over *K* ancestry labels because the data used for the classification task has uniformly distributed ancestry labels. Therefore:

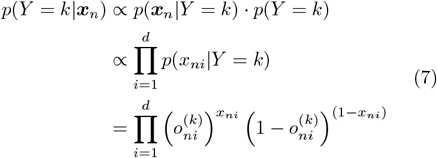

Given that *d* represents a relatively large number of dimensions, numerical computation can lead to the entire expression collapsing to zero. This effect is due to two factors. First, if at least one SNP position is reconstructed incorrectly, i.e. the true value of the SNP position is 0 but the VAE returns 1 or vice versa, the expression becomes zero. Second, when many positions have extremely small values due to numerical precision limitations, the product of these values becomes effectively zero. To address these issues and improve numerical stability, it is common practice operate in the space of log-probabilities. We therefore apply logarithms to Equation 7:

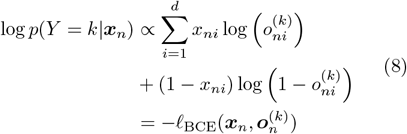

Selecting the ancestry label that maximizes the posterior *p*(*Y* = *k* | ***x***_*n*_) is equivalent to the minimizing the BCE loss:

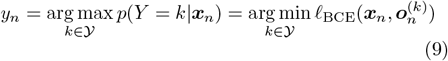

To address the issues associated with the logarithm of zero and the potential misclassification of SNP positions, some form of smoothing or clamping is necessary. One way to implement a form of *Laplace smoothing* is by adjusting the sigmoid temperature, which makes the Bernoulli probabilities 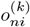 less sharp and less likely to be clamped to zero quickly. However, even with this approach, there can still be issues with zeroing.

To mitigate this, we employ the trick used in the PyTorch implementation of BCE loss [54]. This involves clamping the probabilities to a certain value if they go beyond a specified threshold. Specifically, we clamp *p*(*Y* = *k*|***x***_*n*_) to *e*^−100^. In terms of logarithms, this is equivalent to clamping log *p*(*Y* = *k*|***x***_*n*_) to −100. This approach ensures that the loss remains finite and avoids the issues associated with infinite values due to the logarithm of zero.

Another important conclusion is that the Bernoulli likelihood maximization problem can be reduced to the minimization of the *L*_1_ discrepancy between the input and the output of the VAE: let *𝓁*_1_(***x, o***^(*k*)^) =||***x***−***o***^(*k*)^||_1_ be the *L*_1_ discrepancy between the SNP array ***x*** and its Bernoulli probabilities by VAE conditioned on *k*-th ancestry. And let *b*(***x, o***^(*k*)^) be the multivariate i.i.d. Bernoulli likelihood function. Then, for each *i*-th element in ***x***:

1. 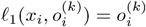 and 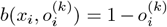 if x_i_ = 0.
2. 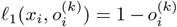 and 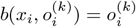 if x_i_ =1.

From what follows the expression:

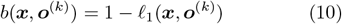

Taking logarithm on Equation 10 yields log *b*(***x, o***^(*k*)^) = log (1 − *𝓁*_1_(***x, o***^(*k*)^)). Since the Taylor series of log (1 − *z*) about zero is:

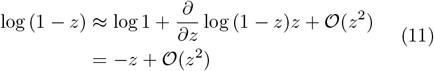

For small *z* all terms of order *z*^2^ are negligible and we can employ the approximation log (1−*z*) ≈ − *z*. Therefore, arg min *𝓁*_BCE_(***x, o***) can be approximated by arg min *𝓁*_1_(***x, o***^(*k*)^). That is, the multivariate i.i.d. Bernoulli likelihood maximization problem can be reduced to the minimization of the *L*_1_ discrepancy between the binary input and output of the VAE.

### Clustering metrics

To quantitatively compare the quality of population groups in both, PCA and VAE spaces, we employ three clustering metrics: the *pseudo F statistic* (also known as Calinski-Harabasz index) [55], the *Davies-Bouldin index* (DBI) [56], and the *silhouette coefficient* (SC) [57]. Pseudo F statistic measures the ratio between the sum of between- and within-cluster dispersion (BSS and WSS, respectively):

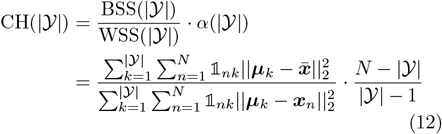

where the number of clusters coincides with the number of populations, denoted as |𝒴|; *N* represents the number of samples in the dataset; 𝟙_*nk*_ serves as an indicator function, determining whether sample ***x***_*n*_, where 1 ≤ *n* ≤ *N*, belongs to ancestry label *k*, with 1 ≤ *k* ≤ |𝒴|; ***x***− represents the mean of the samples, and ***μ***_*k*_ denotes the centroid of the *k*-th ancestry. A higher score indicates that the clusters are more compact and well-separated. Conversely, DBI inversely relates to cluster separation; a smaller DBI value suggests better separation between clusters as it computes the maximal ratio between intra-cluster variance and inter-cluster distance [56]:

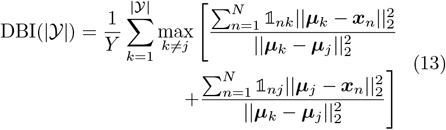

Lastly, SC relates the intra-cluster distance of samples and the nearest-cluster distance. This coefficient is bounded SC(|𝒴|) ∈ [-1, 1]. A value of ∈ [0.71, 1] is an indicator of a strong structure in the data. Values below *<* 0.25 indicate that no substantial structure has been found [57].

## Results

### Dimensionality reduction with VAE

Genotypes can unravel population structure. The identification of genetic clusters can be important when performing GWAS and provides an alternative to self-reported ethnic labels, which are culturally constructed and vary according to the location and individual. A variety of unsupervised dimensionality reduction methods have been explored in the past for such applications, including PCA, MDS, t-SNE, and UMAP. Recently, VAEs have been introduced into population structure visualization [39, 58]. The singular feature of VAEs is that they can represent the population structure as a Gaussian-distributed continuous multi-dimensional representation and as classification probabilities providing flexible and interpretable population descriptors. Besides, latent maps allow for meaningful interpretation of distances between ancestry groups.

We quantitatively assess the quality of the VAE clusters by comparing the clustering performance of PCA and VAE clusters. We use the Pseudo F statistic [55], DBI [56], and SC [57] as clustering metrics. Refer to the **Methods** section for a detailed explanation of these metrics. The results are shown in Table 1.

**Table 1.** Comparison of clustering performance of PCA vs VAE: PCA and VAE parameters have been fitted to human and canine SNP datasets of 839,629 and 198,473 SNP positions, respectively. Clustering metrics have been computed on 6 and 16 ancestry groups, respectively. The latent coordinates of the samples have been standardized.

|  | **Data type:** | Human data | Canine data |
| --- | --- | --- | --- |
| Pseudo F statistic | PCA | 50315.56 | 151.89 |
|  | VAE | **56409.29** | **276.03** |
| Silhouette Coefficient | PCA | 0.69 | **0.12** |
|  | VAE | **0.77** | 0.07 |
| Davies-Bouldin index | PCA | 0.48 | 3.87 |
|  | VAE | **0.29** | **3.39** |

Visually, the VAE clusters still preserve the geographic vicinity of adjacent human populations and, additionally, discriminate more than PCA some subpopulations within each cluster, as it can be clearly observed in the case of African (AFR) and Native American (AMR) populations in Figure 1.b. Therefore, VAE projections to the two-dimensional space allow for a more insightful and fine-grained exploratory analysis. As an example, having a closer look to the aforementioned ancestry groups, shown in Figure 1.b, observe that PCA is not capable of differentiating their sub-populations. In contrast, VAE clusters in the African population clearly distinguish Mbuti and Biaka sub-populations, which both are pygmy populations from the central African cluster. Another interesting visualization is the projection of canine genotypes to the VAE latent space (Figure 1.a). For instance, the Asian Spitz clade is found closer to the wolves, which suggests their genetic similarity as they were one of the first domesticated canids [59].

**Fig. 1.**
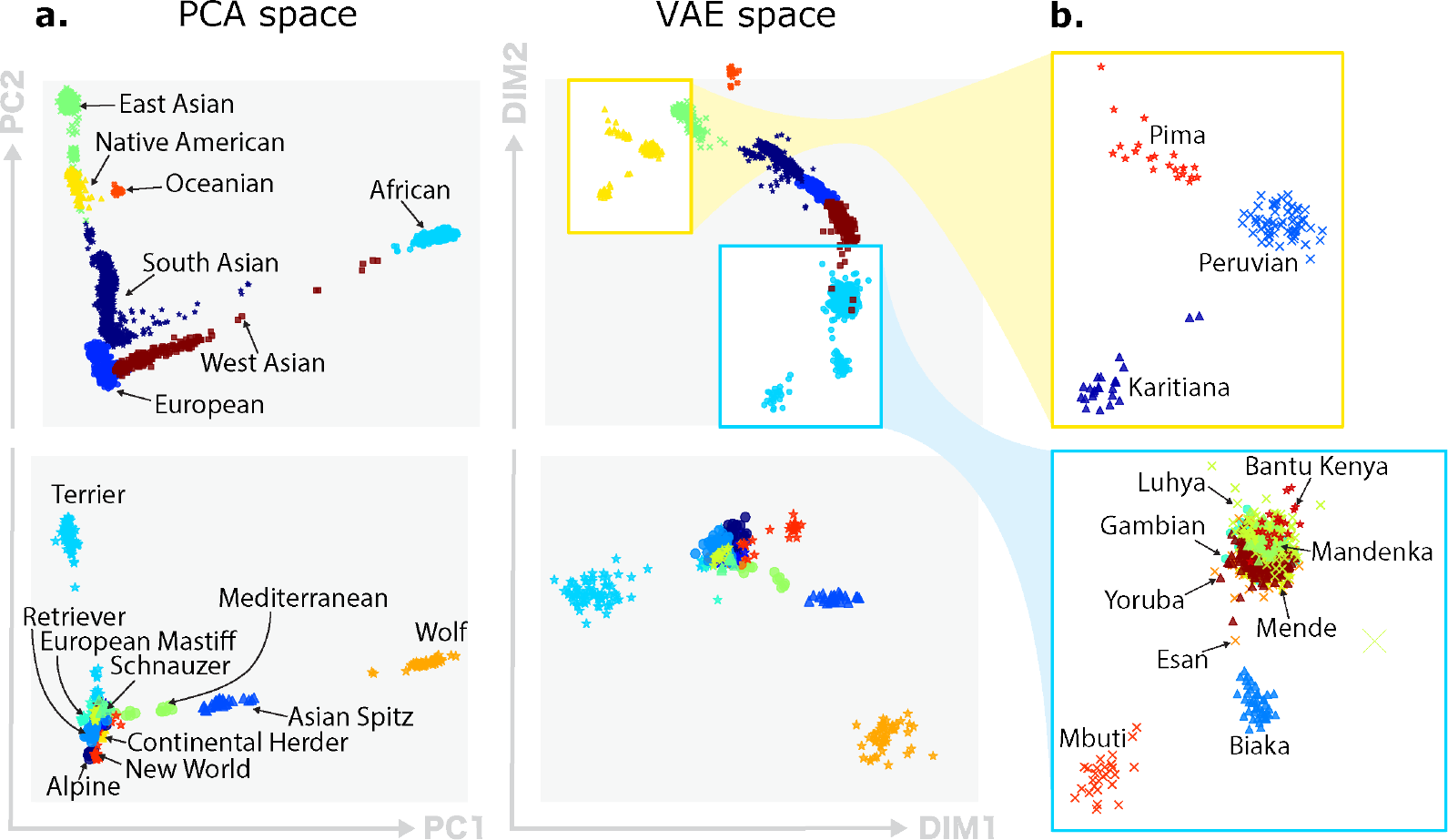
Qualitative comparison of PCA and VAE projections. **(a)** The top row illustrates the projections generated by both PCA and VAE for 4,894 human samples using 839,629 SNPs. The second row displays projections of 489 canine samples using 198,473 SNP positions. **(b)** Focus of VAE projections of Native American subpopulations (in yellow), and African subpopulations (in blue).

### Different objectives for ancestry classification

Individuals that are more closely related ancestry-wise have spatial autocorrelations in their genomic sequences. This phenomenon is translated into population clusters using dimensionality reduction techniques (Figure 1). The most common approaches to address ancestry classification and regression are based on dimensionality reduction and clustering techniques. Genomic sequences from known and unknown origins are jointly analyzed – unknown samples are assigned to the nearest labelled cluster of the feature space (e.g., PCA space). Yet there are some caveats [14, 60]: (1) The results can be nonsensical if the individuals to classify are admixed, i.e. are descendants of individuals from different ancestries, or do not originate from any of the sampled reference populations (*out-of-sample*); (2) Commonly used dimensionality reduction techniques such as PCA do not model LD. Correlations induced by LD violate the inherent assumptions of independency between SNP positions. The cumulative effect of those correlations not only decreases accuracy but can also bias the results, a fact that can be observed in PCA projections of sequential SNP positions compared to SNP positions selected at random (refer to **Appendix** 1 for details). In contrast, VAE, the non-linear counterpart of PCA, is able to model up to a certain degree these induced correlations allowing for less LD bias with a relatively small number of SNP positions as input and consequently, yielding better visualizations. For this task, we have trained VAEs on 10,000 sequential SNP positions from human chromosome 22. The learned representations with VAEs have been assessed based on the population labels we have for each individual. We provide a quantitative evaluation of these learned representations using different classification approaches which are described in detail next.

The simplest idea for classification in the latent space is computing the population centroids and assigning each individual to the nearest one. We refer to this method as the *nearest latent centroid* heuristic. It is a form of nearest-neighbor classification based on the latent features extracted by the VAE. Each centroid ***c***_*k*_, 1 ≤ *k* ≤ |𝒴| is computed as:

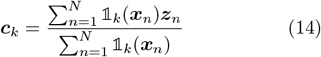

where ***z***_*n*_ = 𝒱_*e*_(***x***_*n*_) and 𝟙_*k*_(·) is an indicator function that denotes membership to ancestry label *k*. For each ***x***_*n*_, we assign the label that minimizes the distance between the latent representation and the corresponding centroid:

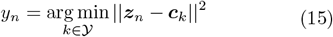

In the classification results presented in this study we set |𝒴|= 7 as we have seven human superpopulations (continental) groups in the provided dataset. We hypothesized that each ancestry-conditioned VAE would better reconstruct the population it belongs to. In the case of Y-VAE, each independent VAE learns to reconstruct better the population on which it has been trained, which is translated into the minimization of the *L*_1_ norm between the input SNP array and the reconstruction. Let us denote the composition of encoder 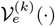, decoder 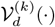, and binarization 𝟙_½_(·) functions, as 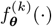 to which we refer to as the VAE model conditioned on *k*-th ancestry with parameters ***θ***. Then, the ancestry of the *n*-th sample:

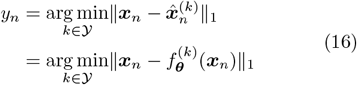

With such conditioning a Bayesian parameter estimation approach can be adopted for ancestry label inference via MAP estimation. Refer to the **Methods** section for the complete mathematical derivation.

The encoder-decoder architecture, when conditioned on *k*-th ancestry, produces a vector of Bernoulli probabilities 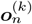, which is subsequently thresholded to generate 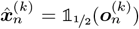. By training the network with the BCE loss we assume that each individual SNP position *x*_*ni*_, 1≤*i*≤*d* follows a Bernoulli distribution. This assumption allows us, in theory, to calculate the Bernoulli likelihood for each SNP position. A MAP estimate of *Y* is the one that maximizes the posterior probability *p*(*Y* = *k* ***x***_*n*_) for a given sample ***x***_*n*_, which is given by:

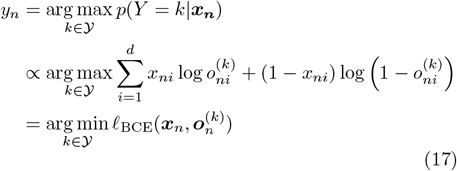

Note that the BCE minimization problem can be approximated by the minimization of the *L*_1_ discrepancy between the input and the output of the VAE. Refer to the **Methods** section for details.

To conclude the classification methods section, we present the results for each method in Table 2. The first two rows compare the *nearest latent centroid* approach in PCA and VAE spaces. Notably, there is a substantial improvement when using the VAE-generated space instead of PCA. The overall test accuracy increases from 74.1% to 85.7%, representing a *>* 15% increase in accuracy. Both C-VAE and *Y* -VAE are evaluated based on two criteria: the minimization of the discrepancy between the input and the reconstruction, and the maximization of the Bernoulli likelihood. Among these, C-VAE, which maximizes the BCE loss, demonstrates the best performance across all populations, achieving an accuracy of 87.1% on the test data. It is worth noting that Oceanian (OCE) and West Asian (WAS) populations exhibit the lowest classification accuracy. Coincidentally, those two populations had the smallest number of founders for simulation. One possible explanation for this phenomenon is that the variability within the simulated samples is insufficient to provide robust generalization for the VAE, necessitating a larger number of founders for improved performance.

**Table 2.** Accuracy of classification methods: *TR* refers to accuracy computed on training data and *TS* on test data, accordingly. The values represent the accuracy in %. Note that regular VAE, C-VAE and Y-VAE have 10, 371, 760; 10, 378, 928, and 72, 602, 320 parameters, respectively.

| Model | Criterion | All |  | EUR |  | EAS |  | AMR |  | SAS |  | AFR |  | OCE |  | WAS |  |
| --- | --- | --- | --- | --- | --- | --- | --- | --- | --- | --- | --- | --- | --- | --- | --- | --- | --- |
|  |  | TR | TS | TR | TS | TR | TS | TR | TS | TR | TS | TR | TS | TR | TS | TR | TS |
| PCA | $\arg \min_k \ z_n - c_k\ _2^2$ | 78.6 | 74.1 | 64.9 | 66.3 | 71.1 | 74.3 | 87.2 | 77.8 | 58.5 | 57.2 | 97.5 | 93.4 | 95.4 | 76.6 | 75.6 | 73.4 |
| VAE |  | 93.2 | 85.7 | 81.7 | 78.6 | 96.9 | 96.3 | 99.5 | 92.1 | 81.6 | 78.5 | 99.3 | 96.8 | 99.4 | 71.7 | 94.0 | 86.1 |
| C-VAE | $\arg \min_k \ x_n - \hat{x}_n^{(k)}\ _1$ | 93.4 | 78.0 | 84.4 | 70.6 | 96.4 | 92.1 | 99.9 | 92.4 | 84.4 | 76.4 | 99.9 | 96.8 | 100 | 62.8 | 88.5 | 54.7 |
| | $\arg \max_k p(Y = k \vert x_n, \theta)$ | 97.5 | **87.1** | 96.4 | **87.1** | 98.5 | **95.5** | 100 | **97.4** | 90.0 | **81.2** | 99.4 | **94.9** | 100 | **79.8** | 98.1 | **73.5** |
| Y-VAE | $\arg \min_k \ x_n - \hat{x}_n^{(k)}\ _1$ | 98.9 | 83.2 | 96.9 | 81.5 | 99.6 | 96.2 | 99.9 | 87.2 | 98.5 | 90.1 | 100 | 98.4 | 100 | 68.6 | 97.5 | 59.9 |
| | $\arg \max_k p(Y = k \vert x_n, \theta)$ | 99.1 | 85.2 | 97.6 | 84.3 | 99.7 | 96.6 | 100 | 90.9 | 98.2 | 88.5 | 100 | 98.2 | 100 | 72.7 | 98.3 | 65.0 |

### Synthetic data generated by VAE

Generating synthetic data with VAE methods is reasonably straightforward. In the initial approach, a regular VAE (without conditioning) has been used to compute the centroids and variances of each population in the latent space, ***μ*** and ***σ***^2^, respectively. With these central points for each cluster, we sample from the isotropic multivariate Gaussian distribution, 𝒩 (***μ, σ***^2^***I***), and decode the resulting latent vector with the VAE decoder 𝒱_*d*_(·). However, this approach does not yield distinct clusters since the distances between the centroids are not sufficiently large to differentiate between populations (Figure 2.a). Based on the insights gained from our simulation experiments, we have recognized that conditioning is essential. The refined simulation algorithm involves sampling a multivariate Gaussian vector, ***z*** ∼ 𝒩 (**0, 1**), and then conditioning the decoder 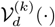 to map this *q*-dimensional latent vector into a *d*-dimensional simulated SNP array 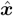 of *k*-th ancestry.

**Fig. 2.**
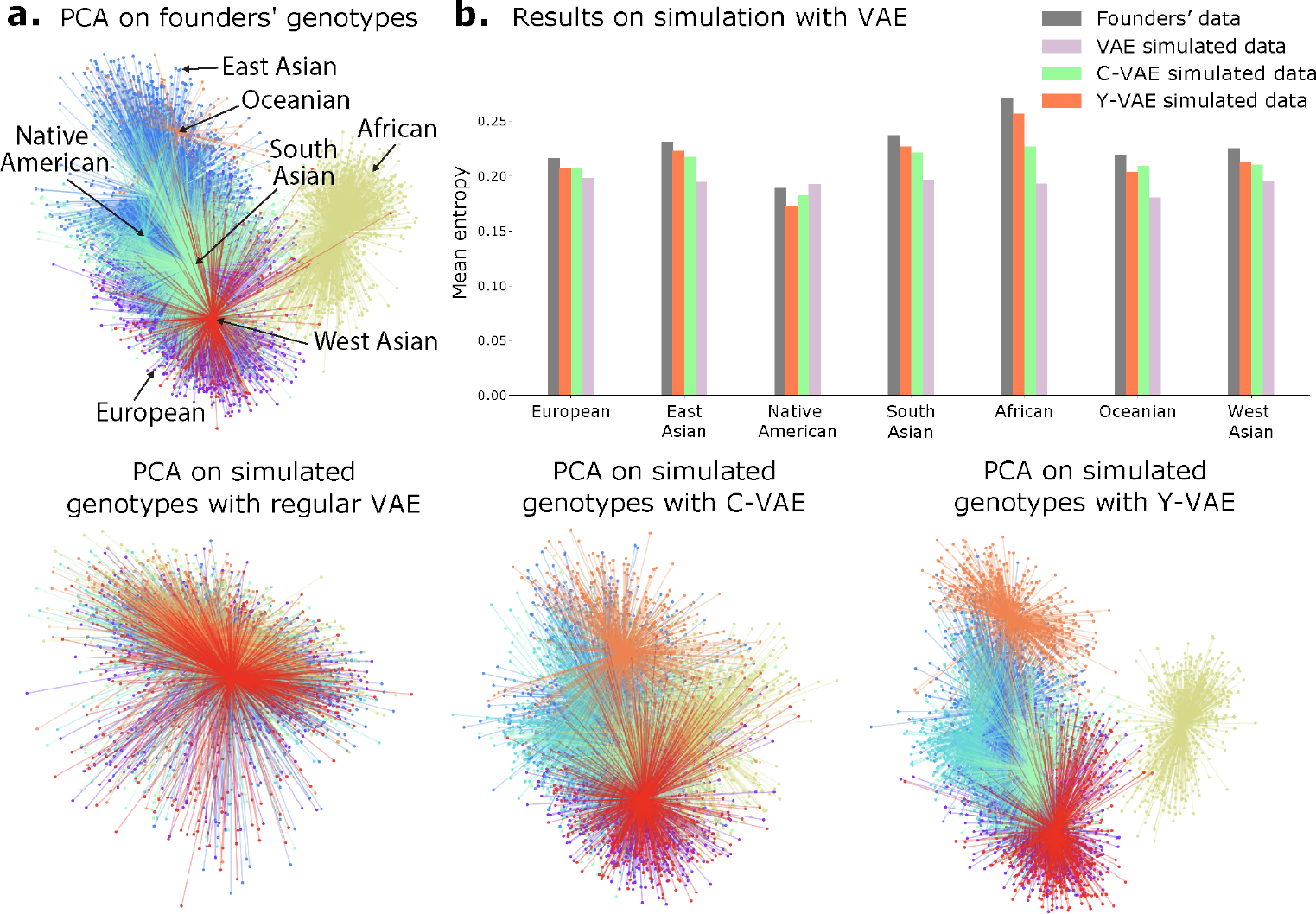
PCA of simulated genotypes and entropy comparison: the number of simulated samples is equal to the number of founders’. **(a)** The top-left plots display the PCA projection of founders’ genotypes. The plots at the bottom line display PCA projections of synthetic genotypes simulated with regular VAE, C-VAE and Y-VAE, in that order. **(b)** The method that best approximates the entropy distribution is Y-VAE. The least effective method is without conditioning, since the entropy is approximately the same across all populations.

In order to quantitatively assess the quality of the simulated individuals, we employ the entropy measure as previously described in [61]. In a genotype array denoted as ***X*** = [*X*_1_, …, *X*_*d*_]^1^, each SNP *X*_*i*_, where 1 ≤ *i* ≤ *d*, is considered a random variable taking values in the Boolean domain B = {0, 1} with probabilities *p*_*ij*_ = *Pr*(*X*_*i*_ = *x*_*j*_), where *p*_*i*_ represents the probability mass function for *X*_*i*_. These probabilities are typically modeled using a Bernoulli distribution. The entropy of *X*_*i*_ is defined as [62]:

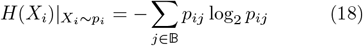

As we anticipate that the simulated samples from a specific ancestry should closely follow the distribution of the corresponding founders (real genotype samples from the simulation pool) comparing the entropy of real and simulated sequences serves as a realiable measure of divergence. As depicted in Figure 2.b, it becomes evident that the best simulation method (Y-VAE) exhibits an entropy distribution that closely resembles that of the founders’. Conversely, the least effective method (unconditional VAE) results in an entropy distribution that is nearly uniform across populations.

### Compression factor with VAE improves over PCA-based approach

Like many natural signals, SNP sequences can be viewed as realizations of a stochastic process. Such data exhibits a particularly high level of redundancy and correlation, owing in part to LD. For that reason, we have conducted a comprehensive entropy analysis to understand the statistical nature of SNP sequences and to estimate the compression bounds.

To estimate the entropies, we leverage the fact that the probability mass function *p*_*i*_ for a Bernoulli distribution is the mean of the random variable. To avoid bias from the unbalancedness of the dataset, we compute the entropy estimates per population by bootstrapping 32 samples from the founders’ pool 50 times and average them. The choice of 32 is significant as it is half of the size of the smallest human superpopulation in the dataset, and the choice of 50 corresponds approximately to the fraction between the size of the largest human continental population divided by 32. The resulting density plots of the entropy rates for human chromosome 22 for each superpopulation are depicted in Figure 3.a. Among these populations, the African population (AFR) presents the highest variability in SNP values and, thus, highest entropy values per SNP. In contrast, the Oceanian (OCE) and Native American (AMR) populations exhibit the lowest values of entropy. These results align with the Out Of Africa (OOA) theory [63], suggesting that populations that migrated out of Africa experienced a reduction in genetic diversity and an increase in LD. Averaging the entropy vectors for each population yields the average SNP entropy per population, as shown in Figure 3.b. These values signal an interesting connection to historical human migrations, as depicted in Figure 3.c. Starting with African individuals, who possess maximum genetic variability, migrations led them to West Asia (WAS), South Asia (SAS), Europe (EUR) and East Asia (EAS). In these regions, populations experienced a noteworthy reduction in average SNP entropy, reflecting a decrease in genetic diversity over time. As time progressed, subsequent migrations brought individuals to Oceania (OCE) and the Americas (AMR), culminating in populations with minimal genetic variability, as evidenced by their lower average SNP entropy. This observation aligns with the notion that genetic diversity tends to decrease as populations migrate further away from their ancestral origins and undergo genetic bottlenecks and founder effects.

**Fig. 3.**
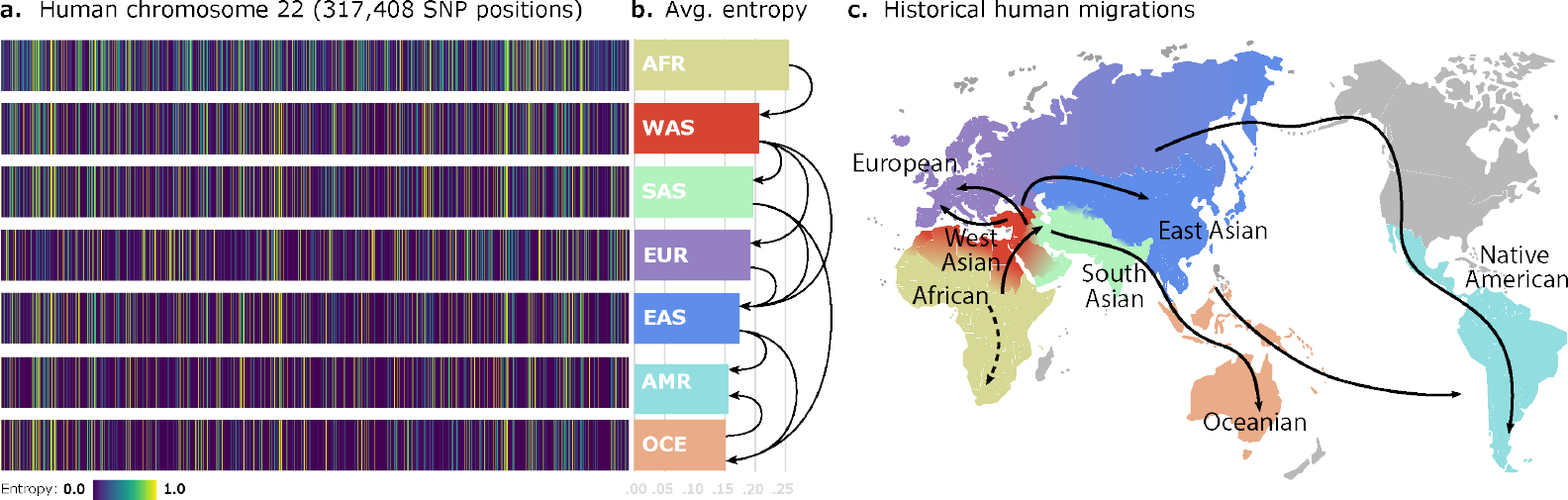
SNP entropy variation between populations. **(a)** We find 317,408 SNP entropies computed for human chromosome 22 for each continental population. Observe that the uncertainty levels for particular SNP positions are different for each population. This is directly related to genetic variability. **(b)** Average of entropy vectors from (a). Those values would correspond to estimated lower bounds of compression for each human population. Arrows represent the migration paths. **(c)** World map showing the main directions of human population migrations.

Based on the VAE architecture, the decoder obtains the reconstruction 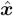, which is a lossy representation of ***x***. To achieve lossless compression, it is necessary to store the residual, which is the element-wise difference between the input and the reconstruction, denoted as 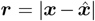, with ***r, x***, 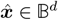. This residual is used for error correction of the reconstruction and with a high-generalization level, a well-trained VAE can produce a sufficiently sparse ***r***. This sparsity can be compressed, in its turn, with another lossless algorithm A. For the sake of simplicity, in our PCA-VAE comparison experiments we use run-length encoding (RLE) for A. We consider a compression execution successful when the inequality from Equation 19 holds:

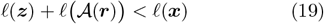

where *𝓁*(·) is the *size function*, which computes the number of digits of the binary representation according to the data type. The latent vector ***z*** is composed of floating-point latent factors, while the reconstructed bits can be stored as Booleans, which, in the best case scenario, should be sparse enough to enable compression with RLE into a smaller integer array.

Table 3 presents the results of human SNP compression for sequences with a length of 10,000, comparing PCA versus VAE. These models were fit to single-ancestry simulated data with 400 generations from founders with 100 individuals in each generation. As observed, VAEs with a bottleneck dimensionality of 32, 64 and 128 latent factors are capable of compressing all populations, including the African population (AFR) which exhibits the highest degree of variability. Notably, European (EUR) and Native American (AMR) ancestries can be compressed to half their original size. In contrast, PCA, being a linear method, struggles to reconstruct effectively from ***z***, resulting in a residual vector ***r*** that is not sufficiently sparse. Consequently, PCA leads to an expansion in size by factors of ×2, ×3, ×4, rather than achieving compression. The VAE takes advantage of the non-linearities in the decoder for improved reconstruction.

**Table 3.** Compression factors of PCA versus VAE: the compression factors are computed as 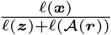 using test data. A compression ratio of 1 corresponds to the identity, values *<* 1 and *>* 1 correspond to compression and expansion, respectively. VAEs with a bottleneck of 32, 64 and 128 latent factors are capable of lossless compression of all human populations. Successful compression is marked in *bold*. |***z***| is the number of latent factors and *α* stands for *learning rate*.

| Models |  |  | Populations |  |  |  |  |  |  |
| --- | --- | --- | --- | --- | --- | --- | --- | --- | --- |
| Type | $\vert z\vert $ | $\alpha$ | European (EUR) | East Asian (EAS) | Native American (AMR) | South Asian (SAS) | African (AFR) | Oceanian (OCE) | West Asian (WAS) |
| PCA | 2 | — | ×0.41 | ×0.38 | ×0.42 | ×0.39 | ×0.33 | ×0.36 | ×0.38 |
| VAE | | $10^{-4}$ | ×0.68 | ×0.64 | ×0.76 | ×0.62 | ×0.53 | ×0.50 | ×0.67 |
| | | $10^{-5}$ | ×0.65 | ×0.61 | ×0.73 | ×0.60 | ×0.52 | ×0.49 | ×0.63 |
| PCA | 4 | — | ×0.41 | ×0.37 | ×0.41 | ×0.38 | ×0.33 | ×0.36 | ×0.38 |
| VAE | | $10^{-4}$ | ×0.77 | ×0.69 | ×0.87 | ×0.68 | ×0.56 | ×0.58 | ×0.71 |
| | | $10^{-5}$ | ×0.73 | ×0.69 | ×0.81 | ×0.66 | ×0.54 | ×0.63 | ×0.68 |
| PCA | 8 | — | ×0.41 | ×0.37 | ×0.41 | ×0.38 | ×0.33 | ×0.36 | ×0.37 |
| VAE | | $10^{-4}$ | ×1.00 | ×0.93 | ×1.17 | ×0.88 | ×0.63 | ×0.68 | ×0.89 |
| | | $10^{-5}$ | ×0.96 | ×0.90 | ×1.08 | ×0.88 | ×0.63 | ×0.74 | ×0.88 |
| PCA | 16 | — | ×0.40 | ×0.36 | ×0.41 | ×0.38 | ×0.32 | ×0.35 | ×0.37 |
| VAE | | $10^{-4}$ | ×1.59 | ×1.39 | ×1.73 | ×1.32 | ×0.84 | ×1.01 | ×1.37 |
| | | $10^{-5}$ | ×1.25 | ×1.12 | ×1.35 | ×1.04 | ×0.70 | ×0.90 | ×1.08 |
| PCA | 32 | — | ×0.39 | ×0.36 | ×0.40 | ×0.37 | ×0.32 | ×0.34 | ×0.36 |
| VAE | | $10^{-4}$ | ×2.00 | ×1.75 | ×2.33 | ×1.69 | ×1.03 | ×1.27 | ×1.72 |
| | | $10^{-5}$ | ×1.67 | ×1.45 | ×1.85 | ×1.39 | ×1.85 | ×1.23 | ×1.28 |
| PCA | 64 | — | ×0.37 | ×0.34 | ×0.38 | ×0.35 | ×0.31 | ×0.33 | ×0.34 |
| VAE | | $10^{-4}$ | ×2.04 | ×1.82 | ×2.27 | ×1.75 | ×1.16 | ×1.47 | ×1.82 |
| | | $10^{-5}$ | ×1.69 | ×1.54 | ×1.96 | ×1.47 | ×0.96 | ×1.30 | ×1.49 |
| PCA | 128 | — | ×0.34 | ×0.32 | ×0.35 | ×0.32 | ×0.29 | ×0.30 | ×0.32 |
| VAE | | $10^{-4}$ | ×1.54 | ×1.45 | ×1.61 | ×1.41 | ×1.06 | ×1.25 | ×1.43 |
| | | $10^{-5}$ | ×1.47 | ×1.37 | ×1.56 | ×1.35 | ×0.97 | ×1.20 | ×1.37 |
| PCA | 256 | — | ×0.30 | ×0.28 | ×0.30 | ×0.28 | ×0.25 | ×0.27 | ×0.28 |
| VAE | | $10^{-4}$ | ×0.97 | ×0.93 | ×1.00 | ×0.93 | ×0.79 | ×0.86 | ×0.93 |
| | | $10^{-5}$ | ×0.94 | ×0.90 | ×0.97 | ×0.89 | ×0.73 | ×0.84 | ×0.90 |
| PCA | 512 | — | ×0.24 | ×0.23 | ×0.24 | ×0.23 | ×0.21 | ×0.22 | ×0.23 |
| VAE | | $10^{-4}$ | ×0.54 | ×0.53 | ×0.55 | ×0.53 | ×0.48 | ×0.50 | ×0.53 |
| | | $10^{-5}$ | ×0.53 | ×0.52 | ×0.54 | ×0.51 | ×0.45 | ×0.49 | ×0.52 |

We also explored compression using a C-VAE. In this setup, the key difference is the inclusion of an additional explanatory variable, denoted as *y*, which encodes the ancestry or breed of the subject. In practical terms, this implies that before compressing a set of SNP arrays, we explicitly condition the encoder based on ancestry or breed. Similarly, during the expansion step, we condition the decoder using that same label. Following our experiments, we opted to continue with the conventional VAE. This decision was based on the slightly worse compression performance observed with C-VAE and its inability to effectively compress the African (AFR) population. A comparison between VAE and C-VAE is provided in Table 4.

**Table 4.** Compression factors of VAE versus C-VAE: the compression factors are computed as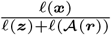 using test data. A compression ratio of 1 corresponds to the identity, values *<* 1 and *>* 1 correspond to compression and expansion, respectively. Successful compression is marked in *bold*.|***z***| is the number of latent factors and *α* stands for *learning rate*.

| Models |  |  | Populations |  |  |  |  |  |  |
| --- | --- | --- | --- | --- | --- | --- | --- | --- | --- |
| Type | $\vert z\vert $ | $\alpha$ | European (EUR) | East Asian (EAS) | Native American (AMR) | South Asian (SAS) | African (AFR) | Oceanian (OCE) | West Asian (WAS) |
| VAE | 2 | $10^{-4}$ | ×0.68 | ×0.64 | ×0.76 | ×0.62 | ×0.53 | ×0.50 | ×0.67 |
| C-VAE |  |  | ×0.57 | ×0.54 | ×0.58 | ×0.54 | ×0.43 | ×0.46 | ×0.55 |
| VAE | 4 | $10^{-4}$ | ×0.77 | ×0.69 | ×0.87 | ×0.68 | ×0.56 | ×0.58 | ×0.71 |
| C-VAE |  |  | ×0.63 | ×0.58 | ×0.63 | ×0.58 | ×0.42 | ×0.51 | ×0.58 |
| VAE | 8 | $10^{-4}$ | ×1.00 | ×0.93 | ×**1.17** | ×0.88 | ×0.63 | ×0.68 | ×0.89 |
| C-VAE |  |  | ×0.78 | ×0.71 | ×0.73 | ×0.69 | ×0.45 | ×0.56 | ×0.70 |
| VAE | 16 | $10^{-4}$ | ×**1.59** | ×**1.39** | ×**1.73** | ×**1.32** | ×0.84 | ×**1.01** | ×**1.37** |
| C-VAE |  |  | ×**1.02** | ×0.77 | ×0.90 | ×0.88 | ×0.50 | ×0.65 | ×0.90 |
| VAE | 32 | $10^{-4}$ | ×**2.00** | ×**1.75** | ×**2.33** | ×**1.69** | ×**1.03** | ×**1.27** | ×**1.72** |
| C-VAE |  |  | ×**1.32** | ×**1.10** | ×**1.23** | ×**1.12** | ×0.59 | ×0.85 | ×**1.14** |
| VAE | 64 | $10^{-4}$ | ×**2.04** | ×**1.82** | ×**2.27** | ×**1.75** | ×**1.16** | ×**1.47** | ×**1.82** |
| C-VAE |  |  | ×**1.32** | ×**1.20** | ×**1.30** | ×**1.18** | ×0.63 | ×0.88 | ×**1.18** |
| VAE | 128 | $10^{-4}$ | ×**1.54** | ×**1.45** | ×**1.61** | ×**1.41** | ×**1.06** | ×**1.25** | ×**1.43** |
| C-VAE |  |  | ×**1.27** | ×**1.20** | ×**1.28** | ×**1.19** | ×0.77 | ×**1.01** | ×**1.20** |
| VAE | 256 | $10^{-4}$ | ×0.97 | ×0.93 | ×1.00 | ×0.93 | ×0.79 | ×0.86 | ×0.93 |
| C-VAE |  |  | ×0.90 | ×0.86 | ×0.91 | ×0.86 | ×0.68 | ×0.78 | ×0.87 |
| VAE | 512 | $10^{-4}$ | ×0.54 | ×0.53 | ×0.55 | ×0.53 | ×0.48 | ×0.50 | ×0.53 |
| C-VAE |  |  | ×0.52 | ×0.50 | ×0.52 | ×0.50 | ×0.44 | ×0.48 | ×0.51 |

### Leveraging autoencoders for lossless SNP compression in practice

The presented results inspire us to push even further the limits of genotype compression. While previously we have used RLE for residuals, in practical production settings, we should transition to more efficient coding methods. Additionally, recognizing that a discrete representation typically consumes less memory than a continuous one, we explore the concept of vector quantization within the VAE framework, known as VQ-VAE [67]. We show that introducing the autoencoder into existing compression pipelines, such as Genozip [64] or Lempel-Ziv codecs [65, 66], can lead to significant improvements compression factors for large SNP datasets.

To discretize the latent representation of ***x***, we employ a quantizer denoted as 𝒬. This quantizer represents ***x*** as a matrix of positive integer indices, which point to a specific set of embeddings, as described in [67]. Formally 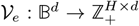, where *H* corresponds to the number of heads in each window-encoder. With a uniform prior, the latent representation is defined by indices pointing to a fixed set of embeddings. We define the codebook size as *K*×*q*, where *K* is the number of embeddings, and *q* signifies the bottleneck dimensionality. We opt for utilizing multi-head encoders with *H* heads. The rationale behind incorporating multiple heads is that the number of potential representations is determined by 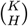. By choosing *H* = 1, we would limit the possible representations to *K* embeddings, which might lead to different inputs mapping to the same quantized latent vector, resulting in non-sparse residuals.

In the context of discrete-space autoencoders, it is important to note that the 𝒬 operator is not differentiable. To address this limitation, we employ the straight-through gradient estimator, which allows for gradient flow during backpropagation. Additionally, within the VQ-objective, we introduce two additional loss terms: (1) the *embedding loss*, which encourages the embeddings to align with the encoder outputs, and (2) the *commitment loss*, which incentivizes the encoder to output ***z*** closer to the embeddings ***e***_*k*_, 1 ≤ *k* ≤ *K*, where *K* is the size of the codebook and *ξ* denotes the stop-gradient operator:

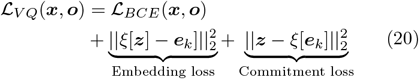

For production purposes, we consider more advanced architectures of autoencoders, specifically, window-based autoencoders (with non-overlapping windows). These autoencoders have two main hyper-parameters: the window size *w* and the bottleneck size *b* for each window. Therefore, the index matrix maintains a fixed size of *H* × (*d/w*). In our window-based architecture an important decision revolved around the selection of values for *w* and *b*. A larger value for *b*, results in more information being compressed into the latent representation, leading to improved reconstruction of ***x*** and consequently a sparser residual. A sparser residual can be better encoded with a coding algorithm *𝒜* because of the longer homogeneous regions of zeros. Concerning the choice of *w*, we have opted for larger window sizes, despite the fact that the intra-window entropy may be higher (which depends on the number of unique sequences in a window). This choice has been motivated by the fact that, even though the intra-window entropy might be higher, the overall sum of expected uncertainty is reduced because we have fewer windows in total.

The window-based encoder 𝒱_*e*_ takes as input a SNP sequence ***x*** of a specific length *d* and compresses it to the dimensionality of the bottleneck *q* = *d/w · b*. In this manner, we obtain the vector of latent factors ***z***, which in its turn is quantized with 𝒬. Next, the decoder 𝒱_*d*_ reconstructs the input with some errors, 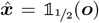, where ***o*** represents the network’s output. As before, storing the residual 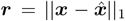, allows for a loss-less compression of ***x***, as the reconstruction errors can be corrected using the residual.With a sufficiently large *q*, the residual ***r*** becomes sparse, making it amenable to compression. This sparsity can be compressed further with a bit stream coding lossless algorithm, along with the latent representation ***z***. The celebrated Lempel-Ziv algorithm belongs to the class of universal compression schemes and in our experiments we use its variations in the role of A.

In this case, we consider a compression execution successful when the Equation 21 holds (note the difference with Equation 19):

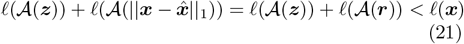

In the proposed method (Figure 4), the window-based VQ-VAE autoencoder is composed of three hidden layers. Each fully connected layer is followed by a batch normalization layer [50]. The activation preceding the bottleneck is transformed by means of the hyperbolic tangent function, to a range which is useful for quantization in the context of discrete latent spaces. The activation at the output of the network is a sigmoid, which converts the activations into Bernoulli probabilities. All the other activation units in intermediate layers are ReLUs. At the beginning of the training process, all the weights and biases are initialized with Xavier initialization [68]. We employ the Adam optimizer [69] with the best performing learning rate of *α* = 0.025 and a weight decay of *γ* = 0.01. A scheduler has been set to reduce the learning rate by a factor of *γ* = 0.1 in the event of learning stagnation. Further, a dropout [70] of 50% has been introduced in all layers because it offers a better generalization providing larger compression factors.

**Fig. 4.**
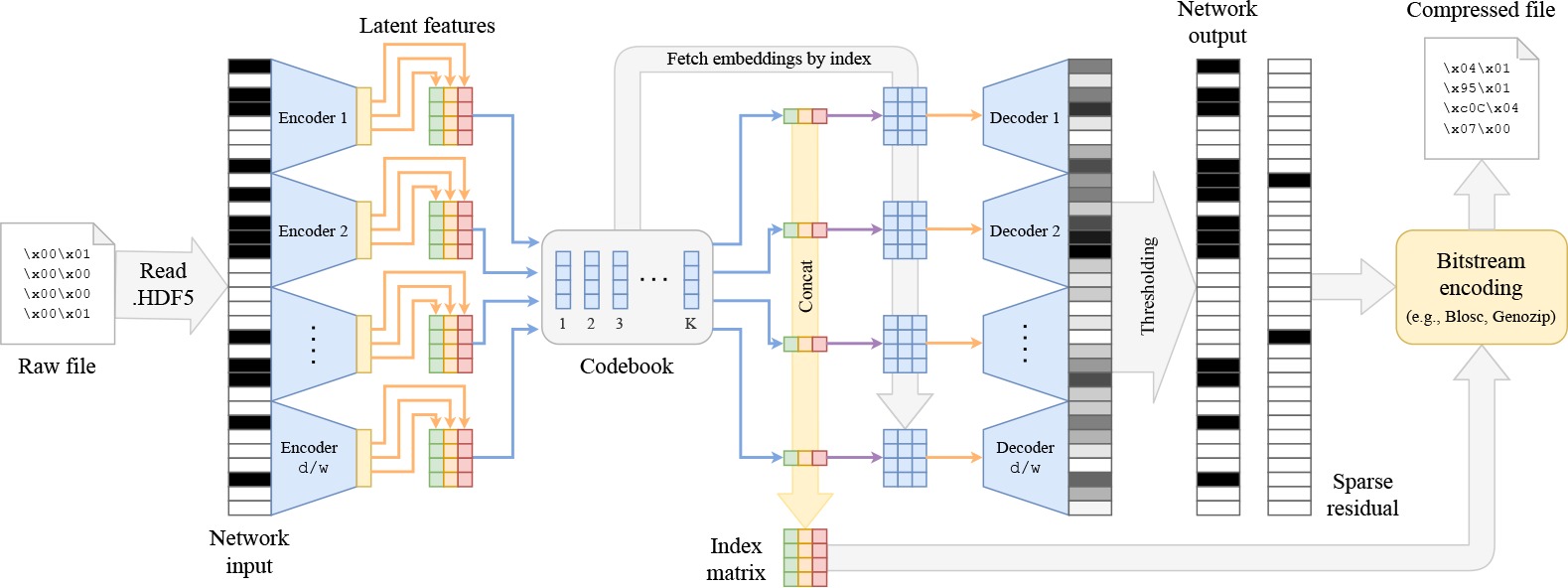
**Proposed VQ-VAE architecture for genotype compression**: the window-based VQ-VAE autoencoder processes an input SNP sequence ***x*** and encodes with 𝒱_*e*_(·) into *H* bottleneck representations (*H* is the number of heads in the encoder). The quantizer 𝒬 substitutes the bottleneck representations by the closest codebook embeddings. Finally, the latent representation can be encoded as an integer index matrix. For the decoding step, codebook embeddings are fetched according to the indices of the index matrix and decoded as usual with the window-based autoencoder. The output is thresholded to obtain the reconstruction. The difference of the input with the reconstruction yields the residual ***r*** which, together with the index matrix, can be integrated in any bitstream-coding-based compression pipeline, such as [64–66].

We benchmark the performance of our autoencoder + bit-stream coding compression strategy against several compression methods, namely: Gzip (general-purpose), ZPAQ [24] (for text), Zstandard [65, 66] (general-purpose) and Genozip [64] (general-purpose optimized for genomic data). We evaluate the compression on three different test sets, each containing 11,772 simulated individuals generated using Wright-Fisher simulation. These test sets consisted of HDF5 files with different numbers of SNPs: 10,000 SNPs from human chromosome 22, and the entirety of human chromosome 22 (approximately 315,000 SNPs), and 80,000 SNPs from human chromosome 20. The results of this benchmark are summarized in Table 5, highlighting the advantages of incorporating autoencoders within compression pipelines.

**Table 5.**
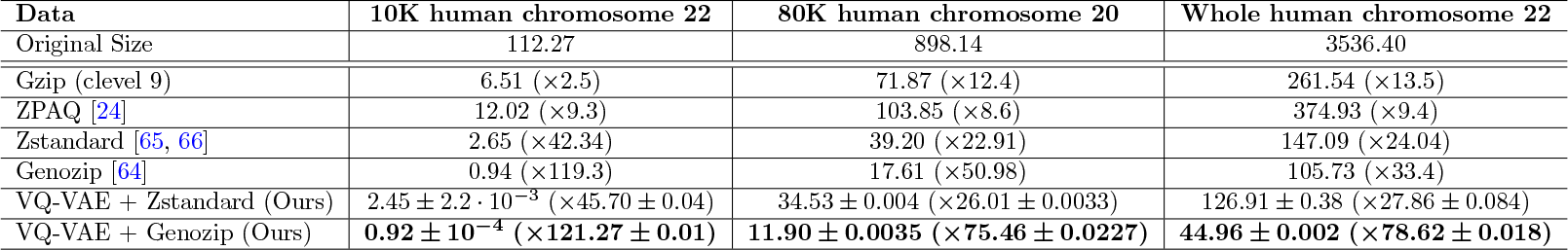
Compression benchmark for SNP sequences. The file size in MB is compared between methods with its compression factor in parenthesis.

| Data | 10K human chromosome 22 | 80K human chromosome 20 | Whole human chromosome 22 |
| --- | --- | --- | --- |
| Original Size | 112.27 | 898.14 | 3536.40 |
| Gzip (clevel 9) | 6.51 ($\times 2.5$) | 71.87 ($\times 12.4$) | 261.54 ($\times 13.5$) |
| ZPAQ [24] | 12.02 ($\times 9.3$) | 103.85 ($\times 8.6$) | 374.93 ($\times 9.4$) |
| Zstandard [65, 66] | 2.65 ($\times 42.34$) | 39.20 ($\times 22.91$) | 147.09 ($\times 24.04$) |
| Genozip [64] | 0.94 ($\times 119.3$) | 17.61 ($\times 50.98$) | 105.73 ($\times 33.4$) |
| VQ-VAE + Zstandard (Ours) | $2.45 \pm 2.2 \cdot 10^{-3}$ ($\times 45.70 \pm 0.04$) | $34.53 \pm 0.004$ ($\times 26.01 \pm 0.0033$) | $126.91 \pm 0.38$ ($\times 27.86 \pm 0.084$) |
| VQ-VAE + Genozip (Ours) | **$0.92 \pm 10^{-4}$ ($\times 121.27 \pm 0.01$)** | **$11.90 \pm 0.0035$ ($\times 75.46 \pm 0.0227$)** | **$44.96 \pm 0.002$ ($\times 78.62 \pm 0.018$)** |

### Imputation with VAE

DNA sequencing data often has missing SNP positions. We show that VAEs can be used to impute the missing values. For this task, we employ a different encoding scheme for the SNP data: we represent the common variant as “−1”, the alternative variant as “1”, and missing values as “0”. We apply a transformation to the input ***x***, such that ***x***^′^ = 2***x*** − **1**, *x*_*i*_ ∈ {0, 1}, and mask missing positions as zeros, ensuring that 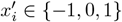 . The VAE attempts to position the missing values either to the negative side (inferring that it belongs to a common variant) or the positive one (imputing it with an alternative variant). Previous imputation works [28] have dealt with different types of missing values, such as *missing completely at random* (MCAR), which we use in our work, and *missing not at random* (MNAR). The MCAR strategy randomly sets any position of the SNP sequence to missing, while in MNAR, missing values are set in specific regions, such as certain genes with particular characteristics. In our experiments we use MCAR with proportions of missing values ranging from 10% to 75%, for 10,000 SNP positions of human chromosome 22. The results are presented in Table 6. We compare the denoising VAE imputation approach with a constant filling method using the common variant. Our results show that the reconstruction accuracy is much better with VAE-based imputation as it learns to impute missing values based on the values of nearby positions.

**Table 6.**
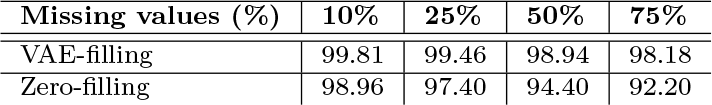
Imputation with VAE compared to constant zero-filling. – the values represent the reconstruction accuracy.

## Discussion

We have demonstrated the power of VAEs applied to population genetics, providing promising performance in a variety of applications. We have conducted both qualitative and quantitative assessments of the quality of VAE clusters on human and canine SNP datasets, reinforcing the benefits of VAE for dimensionality reduction in a population genetics context. We have benchmarked VAE-based global ancestry classification versus PCA-based classification and proved that the non-linear approach works better. Because of the introduced non-linearities, the method is less sensitive to correlations of SNPs due to LD, resulting in an increased ability to capture complex population structure and represent relatively good ancestry differentiation. In contrast to previous work for genotype simulation with VAEs [39], we have used VAE conditioning and found that it is essential in order to achieve high-quality simulated SNP sequences. VAE simulation provides an efficient method for SNP data simulation; we have conducted an exhaustive entropy study on SNP data and reaffirmed the hypothesized migration paths [63] and phylogenetic relationships of the population groups, along with the theoretical compression bounds based on the statistical nature of SNP data. Finally, we have developed a novel, VAE-based, lossless compression system tailored for SNP data.

Undoubtedly, the entropy analysis has proven to be a valuable tool for uncovering and tracing the relationships between population groups. Further exploration of entropy and mutual information at a finer-grained level, including analyses involving different species beyond humans and canids, could yield interesting insights into genetic diversity and population dynamics. The combination of VAEs with entropy analysis, compression techniques, together with genotype classification, simulation, and imputation methods, holds promise for further advancements in our understanding of genetic diversity, ancestry, and evolution.

## Acknowledgements

We thank Alan Aw, Jerry Lin, and Ayesha Bajwa for valuable feedback. In particular, we wish to acknowledge Ayesha for her exceptional assistance in meticulously proofreading the manuscript.

## Abbreviations

ANN: artificial neural network;
VAE: variational autoencoder;
GWAS: genome-wide association study;
SNP: single nucleotide polymorphism;
MLP: multilayer perceptron;
LAI: local ancestry inference;
PPM: prediction by partial matching;
GRU: gated recurrent unit;
LSTM: long short-term memory;
LD: linkage disequilibrium;
GAN: generative adversarial network;
RBM: restricted boltzmann machine;
ReLU: rectified linear unit;
GELU: gaussian error linear unit;
MAF: minor allele frequency;
BCE: binary cross-entropy;
KL: Kullback-Leibler;
MAP: maximum a posteriori;
PCA: principal component analysis;
DBI: Davies-Bouldin index;
SC: silhouette coefficient;
AFR: African;
EUR: European;
AMR: Native American;
WAS: West Asian;
SAS: South Asian;
OCE: Oceanian;
RLE: run-length encoding;
MCAR: missing completely at random;
MNAR: missing not at random;
VQ-VAE: vector quantized variational autoencoder;

## Availability of data and materials

The samples used in the experiments have been compiled using several public data sources: (1) *The 1000 genomes project* [40] (2,504 samples from 26 human populations); (2) *The Human Genome Diversity Project* [41] (929 samples from 54 human populations; (3) *The Simons Genome Diversity Project* [42] (300 samples from 142 human populations); (4) Genotyping array used by the commercial brand Embark [46] (722 canine samples from 144 dog breeds). The source code is publicly available at https://github.com/AI-sandbox/aegen.

## Author contributions

M.G. performed the research and wrote the code base. M.G., D.M.M., X.G.N., and A.G.I. interpreted the results. M.G. and D.M.M. wrote the manuscript.

## Author information

^1^Department of Biomedical Data Science, Stanford University School of Medicine, Palo Alto, CA, USA. ^2^ Department of Signal and Theory Communications, Universitat Politècnica de Catalunya, Barcelona, Spain. ^3^Berkeley AI Research Laboratory (BAIR), Department of Electrical Engineering and Computer Science, University of California at Berkeley, Berkeley, CA, USA. ^4^Institute for Computational and Mathematical Engineering, Stanford University School of Engineering, Palo Alto, CA, USA.

## Appendix

### Relevance of this work for precision medicine

*Pharmacogenetics*, the study of the response of individuals to drug therapy based on their genome, is a key aspect in precision medicine applications. While external factors, such as diet or environment, can have influence on medication response, genomic information plays an important role, with different individuals having different responses to the same drug. Precision medicine aims to maximize the efficacy of drugs and mitigate the risks of side effects by mapping the right drug and the right dose to each individual with the help of genetic data [6]. Because the genome determines the expression of all the organism’s enzymes, including the ones that metabolize drugs, a drug response of an individual is directly dependant on the genetically determined concentration-response relationships of the enzymes. *Pharmacokinetics* (how an organism affects a drug) and *pharmacodynamics* (how a drug affects an organism) can have significant differences at both individual and population-level [71]. To illustrate, in the US, as there is a great ethnic diversity in the overall population, some drug labels include clinical trials’ data on self-identified white and non-white subpopulations because of the possible ethnic differences in drug response. Therefore, the interethnic differences in drug response have become crucial to establish public health policies, to design and evaluate clinical trials, and to develop, approve, and promote new drugs [6].

### List of human populations in our dataset

Bantu from Kenya (11), Mbuti (13), Mandenka (22), Biaka (22), Mende (85), Esan (99), Luhya (100), Gambian Mandinka (113) and Yoruba (130) from African (AFR) ancestries; Pima (12), Karitiana (12) and Peruvian (31) from Native American (AMR) ancestries; Miao (10), She (10), Uygur (10), Yi (10), Tu (10), Northern Ha (10), Daur (10), Yakut (25), Han (33), Dai Chinese (93), Kinh Vietnamese (99), Han Chinese (103), Southern Han Chinese (105) and Japanese (131) from East Asian (EAS) ancestries; Bergamo Italian (12), Orcadian (15), Adygei (16), Basque (23), Russian (25), French (28), Sardinian (28), British (91), Finnish (99), Spanish (107) and Tuscan (115) from European (EUR) ancestries; Bougainville (11) samples from Oceanian (OCE) ancestry; Hazara (19), Kalash (22), Balochi (24), Burusho (24), Pathan (24), Sindhi (24), Brahui (25), Makrani (25), Bengali (86), Punjabi (96), Sri Lankan (102), Indian Telugu (102) and Gujarati (103) from South Asian (SAS) ancestry, and Mozabite (27), Druze (42), Bedouin (46) and Palestinian (46) from West Asian ancestry.

### List of canine populations in our dataset

Pointer Setter (11), Scent Hound (12), Mediterranean (13), Drover (16), Spaniel (17), Poodle (18), New World (20), Alpine (22), Continental Herder (26), Asian Spitz (29), European Mastiff (29), UK Rural (47), Retriever (51), Wolf (51) and Terrier (115) canids.

---

**Fig. 1.**
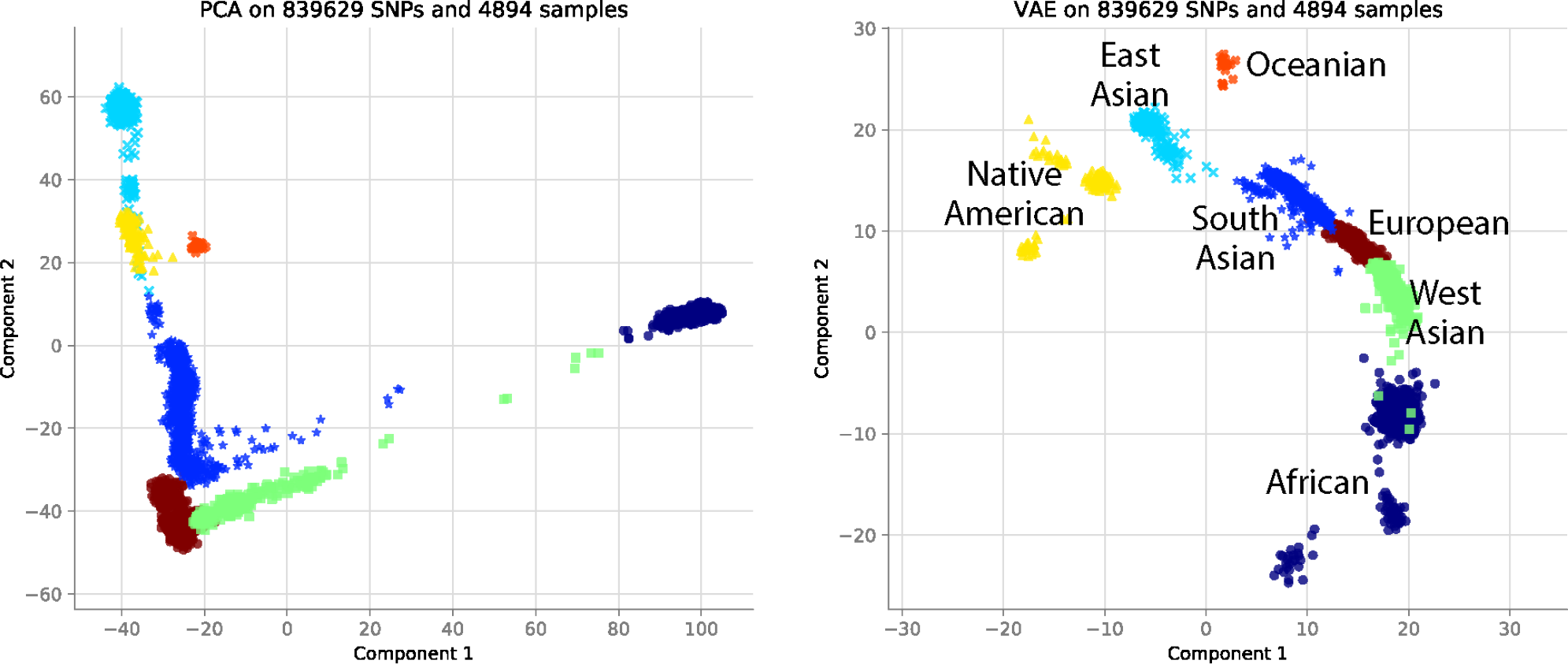
Comparison of PCA and VAE projections of all human population altogether.

**Fig. 2.**
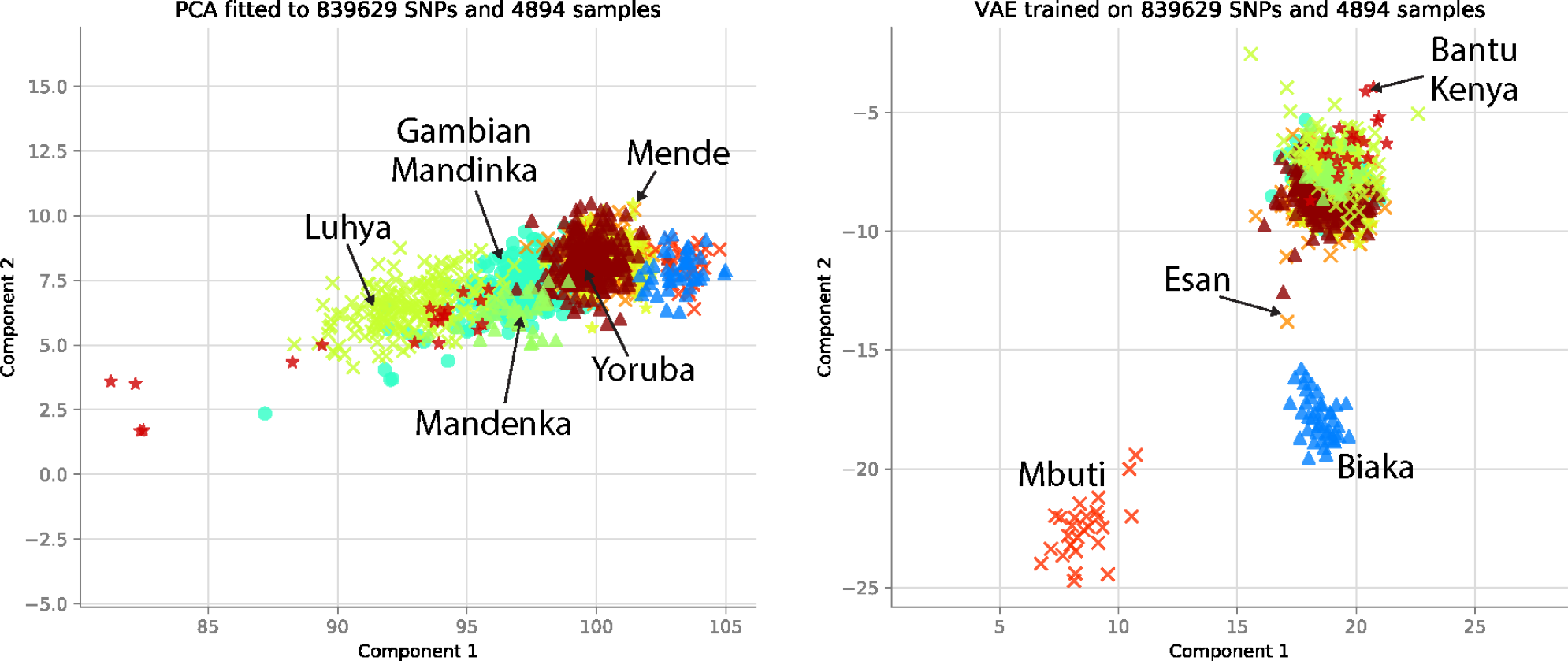
Comparison of PCA and VAE projections of African subpopulations.

**Fig. 3.**
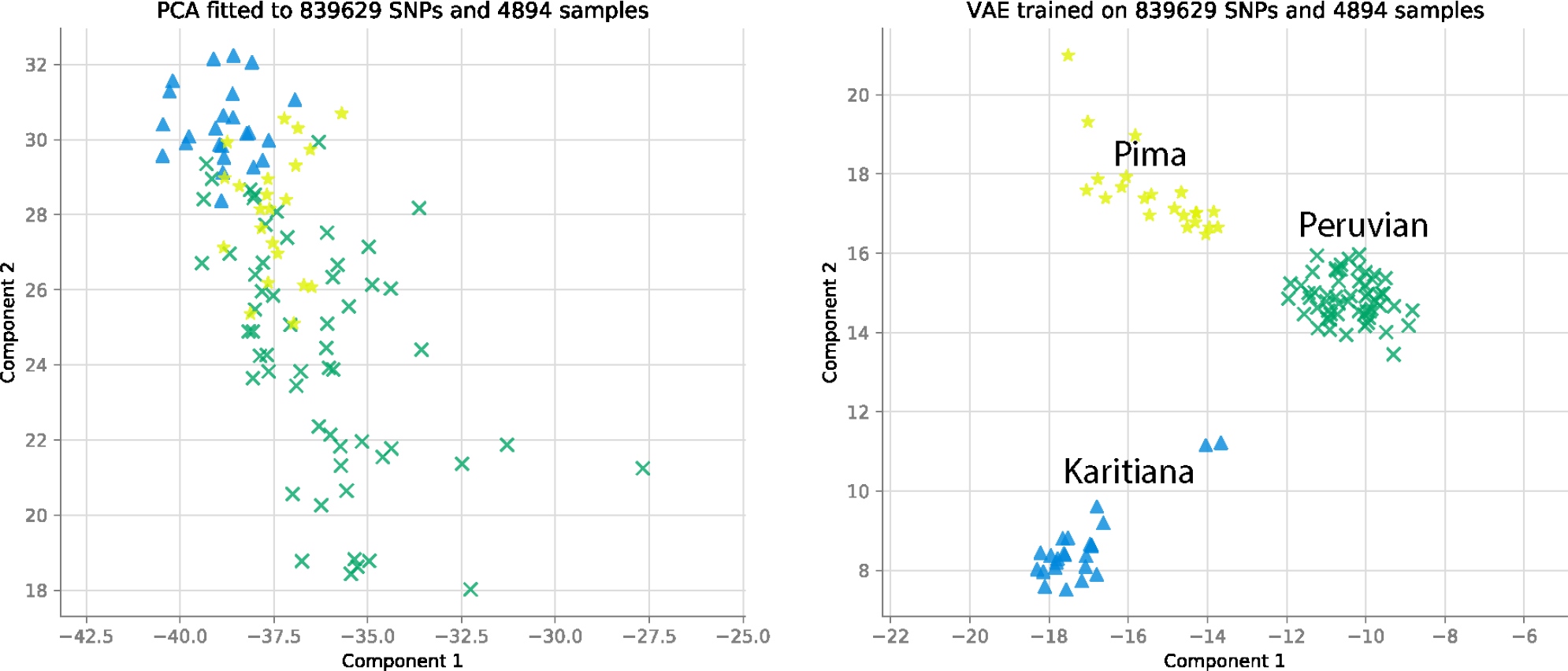
Comparison of PCA and VAE projections of Native American subpopulations.

**Fig. 4.**
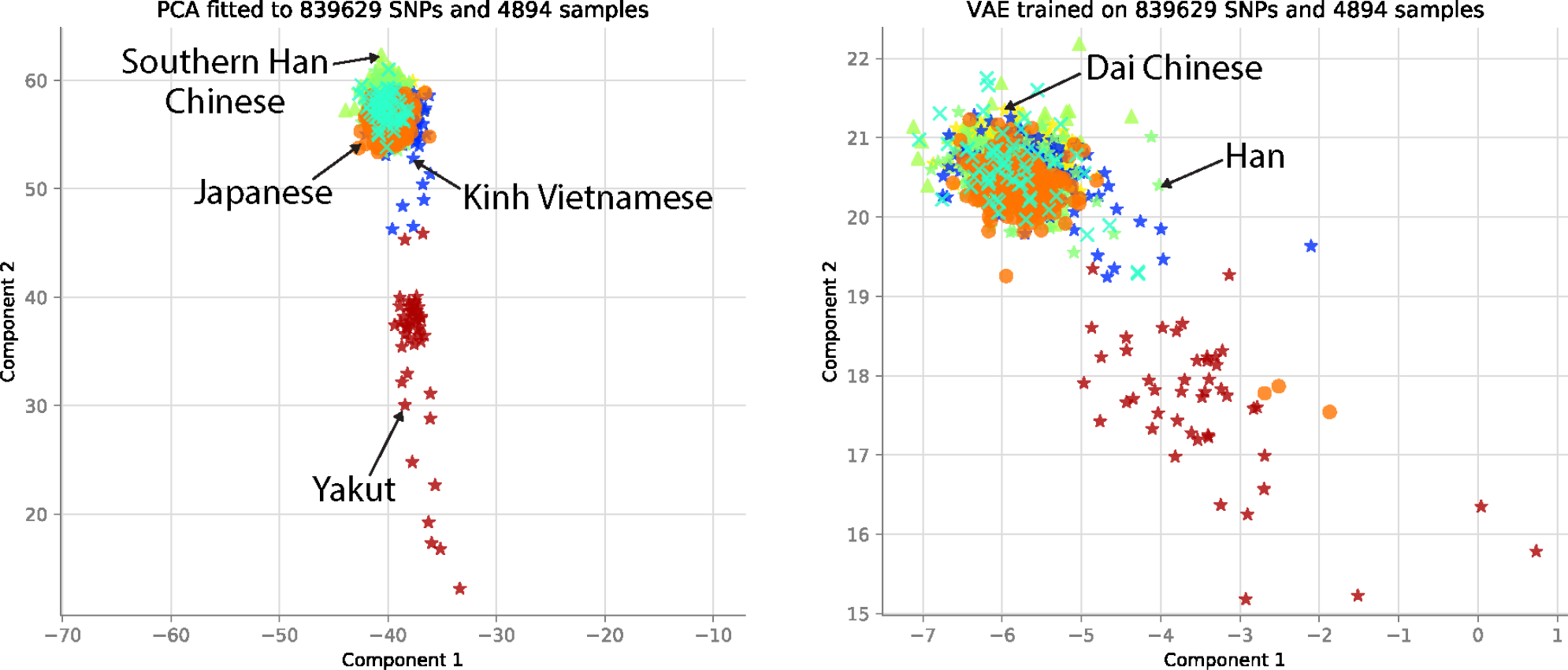
Comparison of PCA and VAE projections of East Asian subpopulations.

**Fig. 5.**
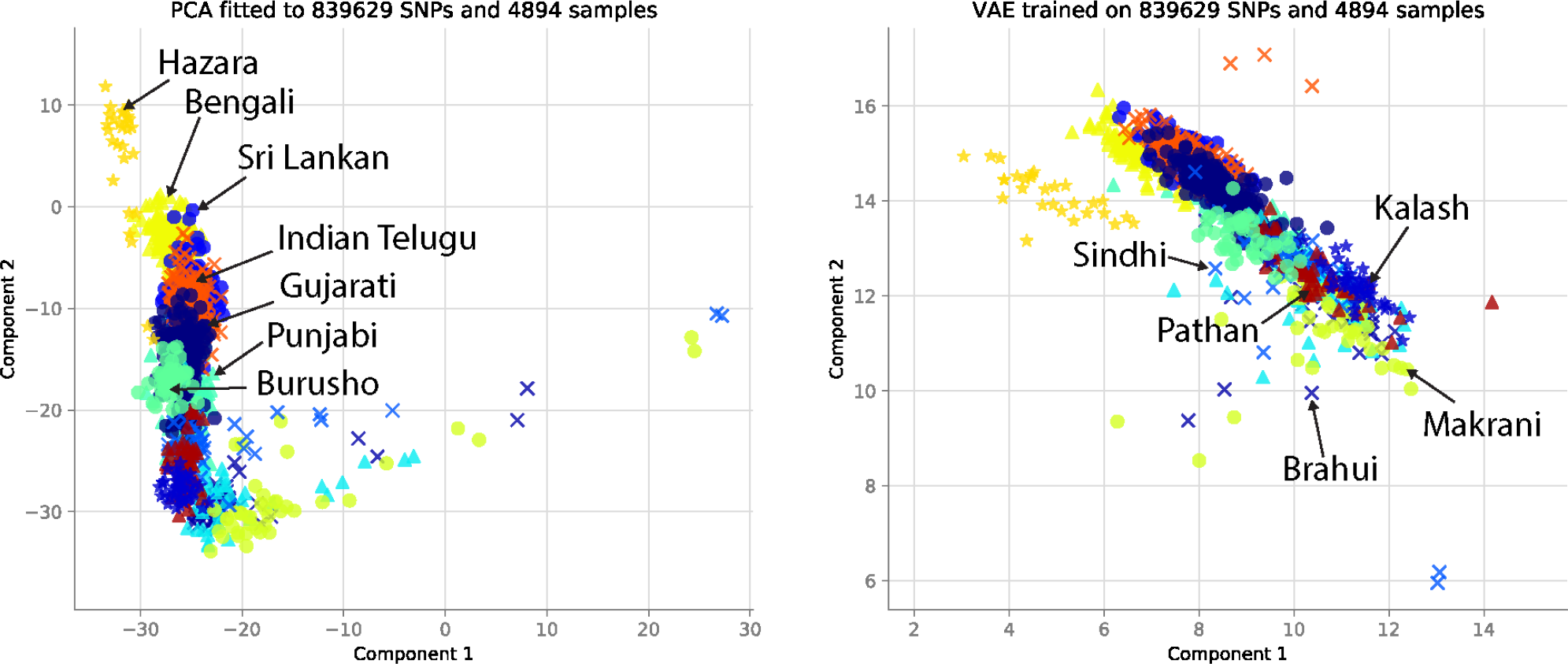
Comparison of PCA and VAE projections of South Asian subpopulations.

**Fig. 6.**
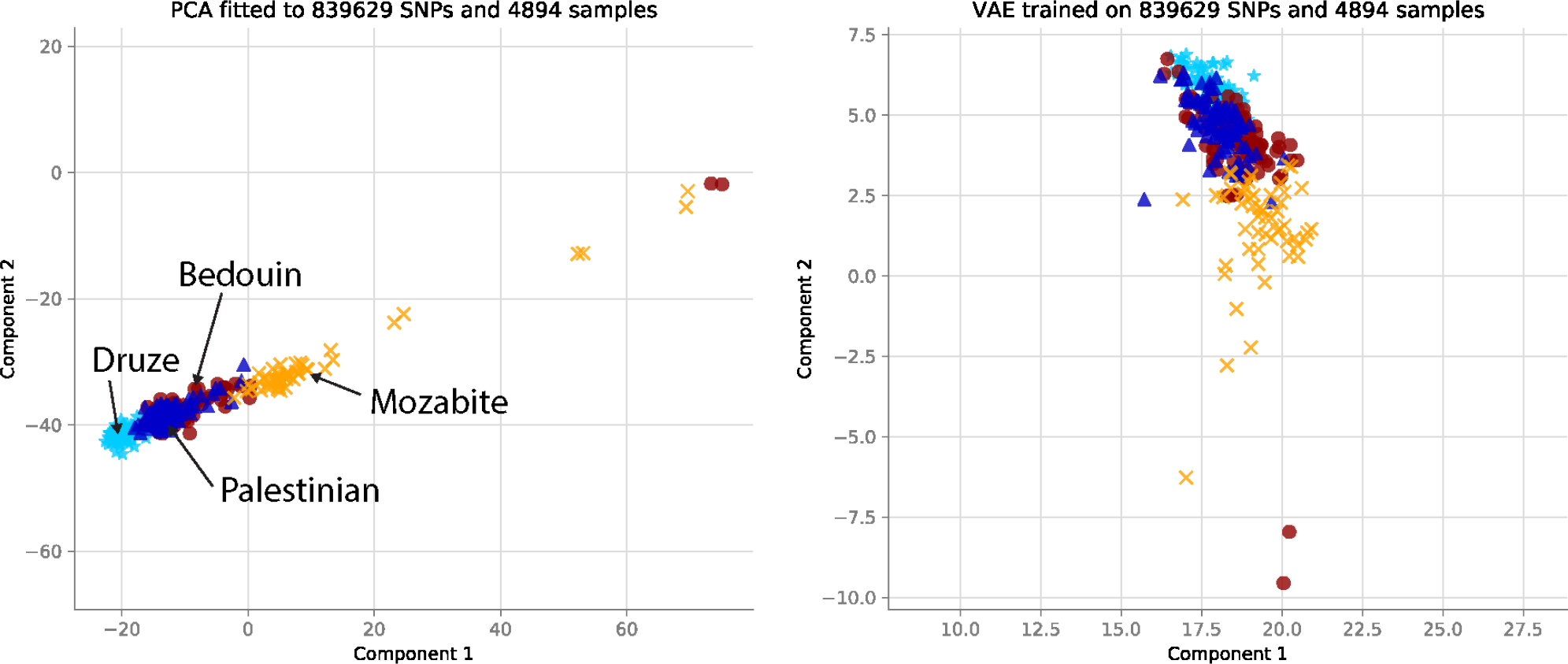
Comparison of PCA and VAE projections of West Asian subpopulations.

**Fig. 7.**
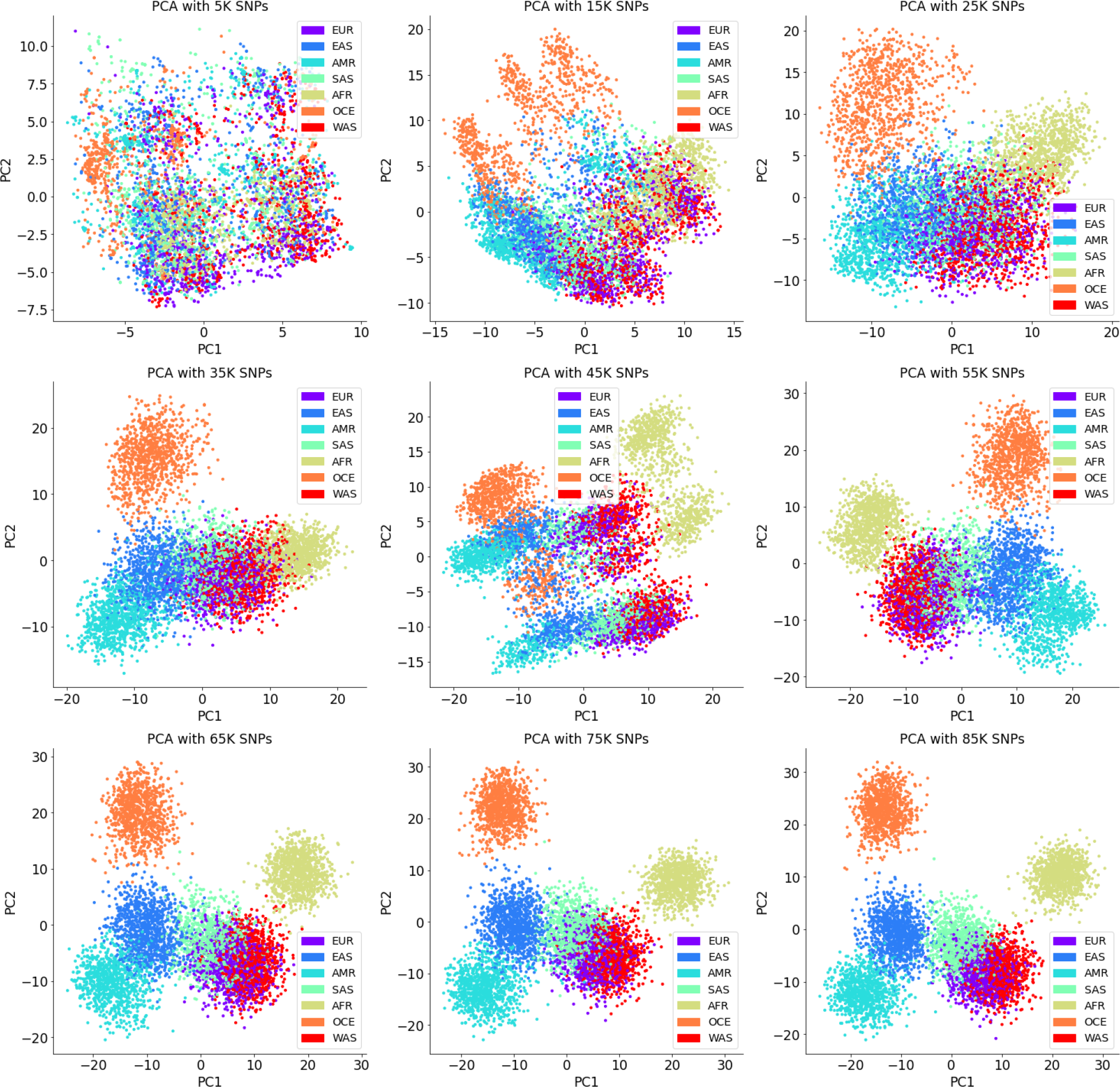
Visual effect of LD on PCA projections. We use sequential SNPs from human chromosome 22 of sizes 5,000; 15,000; 25,000; 35,000; 45,000; 55,000; 65,000; 75,000, and 85,000 SNPs. The *polarization effect* disappears when we use an effectively large subsample of sequential SNPs.

**Fig. 8.**
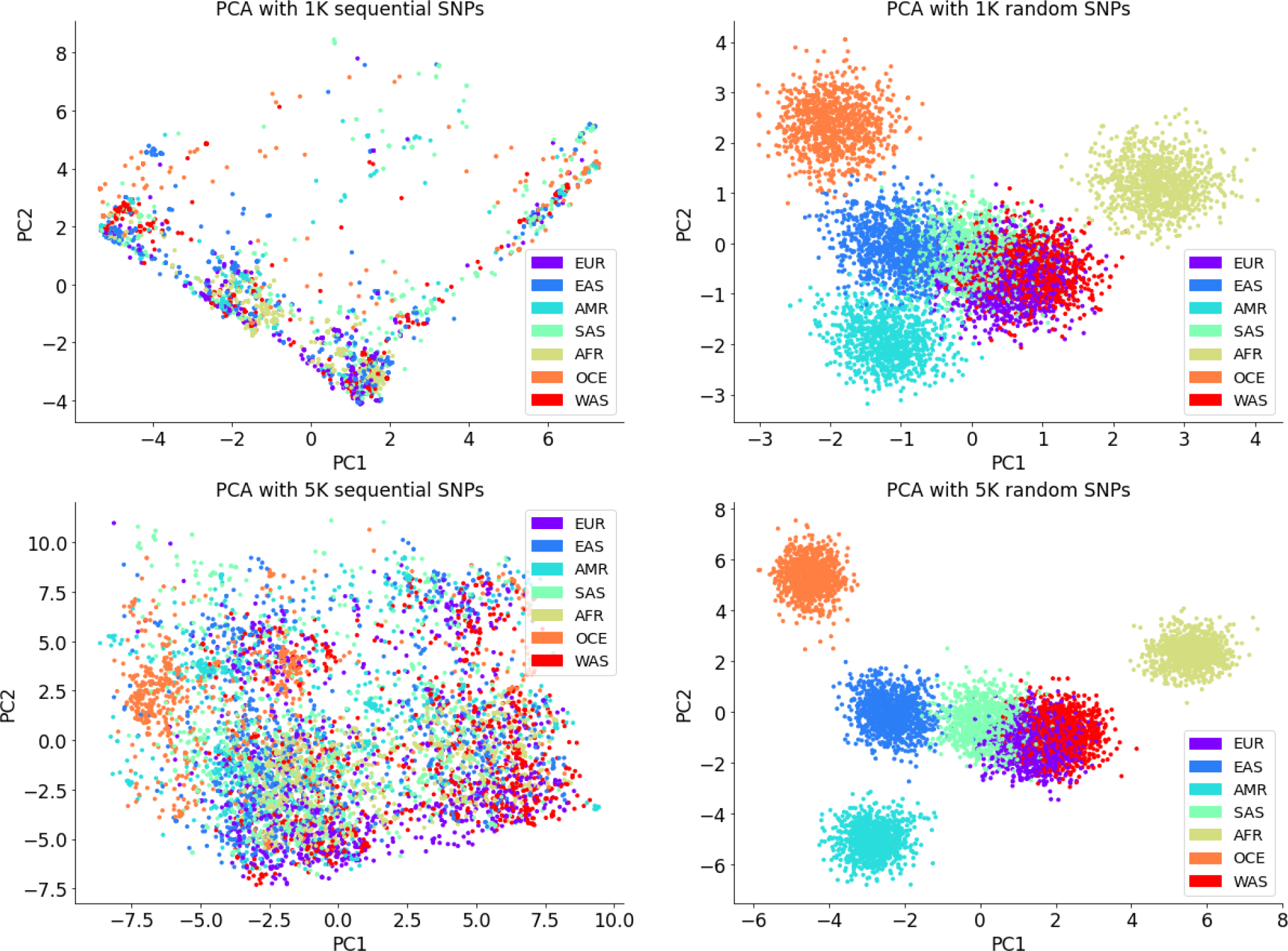
PCA on a subset of random and sequential SNPs. The visual difference is due to LD.

## Footnotes

1 We differentiate between random variables, e.g., ***X***, and samples, e.g., ***x***, using uppercase and lowercase letters, respectively.

## References

[1] Montserrat, D.M., Bustamante, C., Ioannidis, A.: Lai-Net: Local-Ancestry Inference with Neural Networks. In: ICASSP 2020 - 2020 IEEE International Conference on Acoustics, Speech and Signal Processing (ICASSP), pp. 1314–1318. IEEE, Barcelona, Spain (2020). 10.1109/ICASSP40776.2020.9053662 . https://ieeexplore.ieee.org/document/9053662/ accessed 2021-0608

[2] Romero, A., Carrier, P.L., Erraqabi, A., Sylvain, T., Auvolat, A., Dejoie, E., Legault, M.-A., Dubé, M.-P., Hussin, J.G., Bengio, Y.: Diet networks: thin parameters for fat genomics. arXiv preprint arXiv:1611.09340 (2016)

[3] Mantes, A.D., Montserrat, D.M., Bustamante, C., Giró-i-Nieto, X., Ioannidis, A.G.: Neural admixture: rapid population clustering with autoencoders. bioRxiv (2021) 10.1101/2021.06.27.450081

[4] Wojcik, G.L., Graff, M., Nishimura, K.K., Tao, R., Haessler, J., Gignoux, C.R., Highland, H.M., Patel, Y.M., Sorokin, E.P., Avery, C.L., Belbin, G.M., Bien, S.A., Cheng, I., Cullina, S., Hodonsky, C.J., Hu, Y., Huckins, L.M., Jeff, J., Justice, A.E., Kocarnik, J.M., Lim, U., Lin, B.M., Lu, Y., Nelson, S.C., Park, S.-S.L., Poisner, H., Preuss, M.H., Richard, M.A., Schurmann, C., Setiawan, V.W., Sockell, A., Vahi, K., Verbanck, M., Vishnu, A., Walker, R.W., Young, K.L., Zubair, N., Acua-Alonso, V., Ambite, J.L., Barnes, K.C., Boerwinkle, E., Bottinger, E.P., Bustamante, C.D., Caberto, C., Canizales-Quinteros, S., Conomos, M.P., Deelman, E., Do, R., Doheny, K., Fernandez-Rhodes, L., Fornage, M., Hailu, B., Heiss, G., Henn, B.M., Hindorff, L.A., Jackson, R.D., Laurie, C.A., Laurie, C.C., Li, Y., Lin, D.-Y., Moreno-Estrada, A., Nadkarni, G., Norman, P.J., Pooler, L.C., Reiner, A.P., Romm, J., Sabatti, C., Sandoval, K., Sheng, X., Stahl, E.A., Stram, D.O., Thornton, T.A., Wassel, C.L., Wilkens, L.R., Winkler, C.A., Yoneyama, S., Buyske, S., Haiman, C.A., Kooperberg, C., Le Marchand, L., Loos, R.J.F., Matise, T.C., North, K.E., Peters, U., Kenny, E.E., Carlson, C.S.: Genetic analyses of diverse populations improves discovery for complex traits. Nature 570(7762), 514–518 (2019) 10.1038/s41586-019-1310-4 . Accessed 2021-06-15

[5] Mardis, E.R.: A decade’s perspective on DNA sequencing technology. Nature 470(7333), 198– 203 (2011) 10.1038/nature09796 . Accessed 2021-06-15

[6] Shah, R.R., Gaedigk, A.: Precision medicine: does ethnicity information complement genotype-based prescribing decisions? Therapeutic Advances in Drug Safety 9(1), 45–62 (2018) 10.1177/2042098617743393 10.1177/2042098617743393. PMID: 29318005

[7] Roberts, D.E.: Fatal Invention: How Science, Politics, and Big Business Re-create Race in the Twenty-first Century, Faculty Scholarship at Penn Law. 433 (2011)

[8] Xie, H.-G., Kim, R.B., Wood, A.J., Stein, C.M.: Molecular basis of ethnic differences in drug disposition and response. Annual Review of Pharmacology and Toxicology 41(1), 815–850 (2001) https://doi.org/10.1146/annurev.pharmtox.41.1.815 10.1146/annurev.pharmtox.41.1.815. PMID: 11264478

[9] Fujimura, J., Rajagopalan, R.: Different differences: The use of ‘genetic ancestry’ versus race in biomedical human genetic research. Social studies of science 41, 5–30 (2011) 10.2307/40997113

[10] Nelson, S.C., Yu, J.-H., Wagner, J.K., Harrell, T.M., Royal, C.D., Bamshad, M.J.: A content analysis of the views of genetics professionals on race, ancestry, and genetics. AJOB Empirical Bioethics 9(4), 222–234 (2018) 10.1080/23294515.2018.1544177 10.1080/23294515.2018.1544177. PMID: 30608210

[11] Novembre, J., Johnson, T., Bryc, K., Kutalik, Z., Boyko, A.R., Auton, A., Indap, A., King, K.S., Bergmann, S., Nelson, M.R., Stephens, M., Bustamante, C.D.: Genes mirror geography within Europe. Nature 456(7218), 98–101 (2008) 10.1038/nature07331 . Accessed 2021-05-29

[12] Pritchard, J.K., Stephens, M., Donnelly, P.: Inference of population structure using multilocus genotype data. Genetics 155(2), 945–959 (2000) https://www.genetics.org/content/155/2/945.full.pdf

[13] Alexander, D.H., Novembre, J., Lange, K.: Fast model-based estimation of ancestry in unrelated individuals. Genome Research 19(9), 1655– 1664 (2009) 10.1101/gr.094052.109 . Accessed 2021-06-08

[14] Battey, C., Ralph, P.L., Kern, A.D.: Predicting geographic location from genetic variation with deep neural networks. eLife 9, 54507 (2020) 10.7554/eLife.54507

[15] Ali-Khan, S.E., Daar, A.S.: Admixture mapping: from paradigms of race and ethnicity to population history. The HUGO Journal 4(1-4), 23–34 (2010) 10.1007/s11568-010-9145-y . Accessed 2021-06-15

[16] Oriol Sabat, B., Mas Montserrat, D., Giro-i-Nieto, X., Ioannidis, A.G.: Salai-net: species-agnostic local ancestry inference network. Bioinformatics 38(Supplement 2), 27–33 (2022)

[17] Maples, B.K., Gravel, S., Kenny, E.E., Bustamante, C.D.: RFMix: A Discriminative Modeling Approach for Rapid and Robust Local-Ancestry Inference. The American Journal of Human Genetics 93(2), 278–288 (2013) 10.1016/j.ajhg.2013.06.020 . Accessed 2021-06-08

[18] Giancarlo, R., Scaturro, D., Utro, F.: Textual data compression in computational biology: a synopsis. Bioinformatics 25(13), 1575–1586 (2009) 10.1093/bioinformatics/btp117 https://academic.oup.com/bioinformatics/article-pdf/25/13/1575/490639/btp117.pdf

[19] Brandon, M.C., Wallace, D.C., Baldi, P.: Data structures and compression algorithms for genomic sequence data. Bioinformatics 25(14), 1731–1738 (2009) 10.1093/bioinformatics/btp319 . Accessed 2021-05-29

[20] Nalbantoglu, A.U., Russell, D.J., Sayood, K.: Data compression concepts and algorithms and their applications to bioinformatics. Entropy 12(1), 34– 52 (2010) 10.3390/e12010034

[21] Schmidhuber, J., Heil, S.: Sequential neural text compression. IEEE Transactions on Neural Networks 7(1), 142–146 (1996) 10.1109/72.478398

[22] Mahoney, M.V.: Fast text compression with neural networks. In: FLAIRS Conference, pp. 230–234 (2000)

[23] Goyal, M., Tatwawadi, K., Chandak, S., Ochoa, I.: DeepZip: Lossless Data Compression using Recurrent Neural Networks (2018)

[24] Mahoney, M.: Adaptive weighing of context models for lossless data compression. (2005)

[25] Silva, M., Pratas, D., Pinho, A.J.: Efficient DNA sequence compression with neural networks. GigaScience 9(11) (2020) 10.1093/gigascience/giaa119 https://academic.oup.com/gigascience/article-pdf/9/11/giaa119/34251844/giaa119.pdf.giaa119

[26] Wang, R., Bai, Y., Chu, Y.-S., Wang, Z., Wang, Y., Sun, M., Li, J., Zang, T., Wang, Y.: Deepdna: a hybrid convolutional and recurrent neural network for compressing human mitochondrial genomes. In: 2018 IEEE International Conference on Bioinformatics and Biomedicine (BIBM), pp. 270–274 (2018). 10.1109/BIBM.2018.8621140

[27] Absardi, Z.N., Javidan, R.: A fast reference-free genome compression using deep neural networks. In: 2019 Big Data, Knowledge and Control Systems Engineering (BdKCSE), pp. 1–7 (2019). 10.1109/BdKCSE48644.2019.9010661

[28] Qiu, Y.L., Zheng, H., Gevaert, O.: Genomic data imputation with variational auto-encoders. GigaScience 9(8) (2020) 10.1093/gigascience/giaa082

[29] Browning, S.R., Browning, B.L.: Rapid and Accurate Haplotype Phasing and Missing-Data Inference for Whole-Genome Association Studies By Use of Localized Haplotype Clustering. The American Journal of Human Genetics 81(5), 1084–1097 (2007) 10.1086/521987 . Accessed 2021-11-16

[30] Browning, B.L., Zhou, Y., Browning, S.R.: A One-Penny Imputed Genome from Next-Generation Reference Panels. The American Journal of Human Genetics 103(3), 338–348 (2018) 10.1016/j.ajhg.2018.07.015 . Accessed 2021-06-12

[31] Qiu, Y.L., Zheng, H., Gevaert, O.: Genomic data imputation with variational auto-encoders. GigaScience 9(8) (2020) 10.1093/gigascience/giaa082 https://academic.oup.com/gigascience/article-pdf/9/8/giaa082/33571399/giaa082.pdf.giaa082

[32] Chen, J., Shi, X.: Sparse Convolutional Denoising Autoencoders for Genotype Imputation. Genes 10(9), 652 (2019) 10.3390/genes10090652 . Accessed 2021-11-16

[33] Popejoy, A.B., Fullerton, S.: Genomics is failing on diversity. Nature 538, 161–164 (2016)

[34] Maher, B.: Genomics: Bioethics on stage. Nature 524(7565), 289–289 (2015) 10.1038/524289a . Accessed 2021-06-15

[35] Guglielmi, G.: Facing up to injustice in genome science. Nature 568(7752), 290–293 (2019) 10.1038/d41586-019-01166-x . Accessed 2021-06-15

[36] Montserrat, D.M., Bustamante, C., Ioannidis, A.: Class-Conditional VAE-GAN for Local-Ancestry Simulation (2019)

[37] Yelmen, B., Decelle, A., Ongaro, L., Marnetto, D., Tallec, C., Montinaro, F., Furtlehner, C., Pagani, L., Jay, F.: Creating artificial human genomes using generative neural networks. PLoS genetics 17(2), 1009303 (2021)

[38] Perera, M., Montserrat, D.M., Barrabes, M., Geleta, M., Giró-i-Nieto, X., Ioannidis, A.G.: Generative moment matching networks for genotype simulation. In: 2022 44th Annual International Conference of the IEEE Engineering in Medicine & Biology Society (EMBC), pp. 1379–1383 (2022). IEEE

[39] Battey, C.J., Coffing, G.C., Kern, A.D.: Visualizing population structure with variational autoencoders. G3 Genes—Genomes—Genetics 11(1) (2021) 10.1093/g3journal/jkaa036 https://academic.oup.com/g3journal/article-pdf/11/1/jkaa036/38018232/jkaa036.pdf

[40] The 1000 Genomes Project Consortium: A global reference for human genetic variation. Nature 526(7571), 68–74 (2015) 10.1038/nature15393 . Accessed 2021-05-29

[41] Bergströmom, A., McCarthy, S.A., Hui, R., Almarri, M.A., Ayub, Q., Danecek, P., Chen, Y., Felkel, S., Hallast, P., Kamm, J., et al.: Insights into human genetic variation and population history from 929 diverse genomes. Science 367(6484) (2020)

[42] Mallick, S., Li, H., Lipson, M., Mathieson, I., Gymrek, M., Racimo, F., Zhao, M., Chennagiri, N., Nordenfelt, S., Tandon, A., et al.: The simons genome diversity project: 300 genomes from 142 diverse populations. Nature 538(7624), 201–206 (2016)

[43] Gravel, S.: Population genetics models of local ancestry. Genetics 191(2), 607–619 (2012) 10.1534/genetics.112.139808 https://www.genetics.org/content/191/2/607.full.pdf

[44] The International HapMap Consortium: A haplotype map of the human genome. Nature 437(7063), 1299–1320 (2005) 10.1038/nature04226. Accessed 2021-06-14

[45] Plassais, J., Kim, J., Davis, B.W., Karyadi, D.M., Hogan, A.N., Harris, A.C., Decker, B., Parker, H.G., Ostrander, E.A.: Whole genome sequencing of canids reveals genomic regions under selection and variants influencing morphology. Nature Communications 10(1), 1489 (2019) 10.1038/s41467-019-09373-w . Accessed 2021-06-14

[46] Bartusiak, E., Barrabes, M., Rymbekova, A., Gimbernat-Mayol, J., Lopez, C., Barberis, L., Montserrat, D.M., Giro-i-Nieto, X., Ioannidis, A.G.: Predicting dog phenotypes from genotypes (2022)

[47] Tipping, M.E., Bishop, C.M.: Probabilistic principal component analysis. Journal of the Royal Statistical Society. Series B (Statistical Methodology) 61(3), 611–622 (1999)

[48] Hinton, G.E., Salakhutdinov, R.: Reducing the dimensionality of data with neural networks. Science 313, 504–507 (2006)

[49] Kingma, D.P., Welling, M.: Auto-Encoding Variational Bayes (2014)

[50] Ioffe, S., Szegedy, C.: Batch Normalization: Accelerating Deep Network Training by Reducing Internal Covariate Shift (2015)

[51] Krizhevsky, A., Sutskever, I., Hinton, G.E.: ImageNet classification with deep convolutional neural networks. Communications of the ACM 60(6), 84–90 (2017) 10.1145/3065386. Accessed 2021-06-14

[52] Hendrycks, D., Gimpel, K.: Gaussian Error Linear Units (GELUs) (2020)

[53] Ma, J., Yarats, D.: Quasi-hyperbolic momentum and Adam for deep learning (2019)

[54] Paszke, A., Gross, S., Massa, F., Lerer, A., Bradbury, J., Chanan, G., Killeen, T., Lin, Z., Gimelshein, N., Antiga, L., Desmaison, A., Kopf, A., Yang, E., DeVito, Z., Raison, M., Tejani, A., Chilamkurthy, S., Steiner, B., Fang, L., Bai, J., Chintala, S.: PyTorch: An Imperative Style, High-Performance Deep Learning Library (2019)

[55] Calinski, T., Harabasz, J.: A dendrite method for cluster analysis. Communications in Statistics 3(1), 1–27 (1974) 10.1080/03610927408827101

[56] Davies, D.L., Bouldin, D.W.: A cluster separation measure. IEEE Transactions on Pattern Analysis and Machine Intelligence PAMI-1(2), 224– 227 (1979) 10.1109/TPAMI.1979.4766909

[57] Kaufman, L., Rousseeuw, P.: Finding Groups in Data: An Introduction to Cluster Analysis, (2009)

[58] Meisner, J., Albrechtsen, A.: Haplotype and population structure inference using neural networks in whole-genome sequencing data. bioRxiv (2020) 10.1101/2020.12.28.424587

[59] Yang, H., Wang, G.-D., Wang, M., Ma, Y., Yin, T., Fan, R., Wu, H., Zhong, L., Irwin, D., Zhai, W., Zhang, Y.: The origin of chow chows in the light of the east asian breeds. BMC Genomics 18 (2017) 10.1186/s12864-017-3525-9

[60] Baran, Y., Quintela, I., Carracedo, A., Pasaniuc, B., Halperin, E.: Enhanced Localization of Genetic Samples through Linkage-Disequilibrium Correction. The American Journal of Human Genetics 92(6), 882–894 (2013) 10.1016/j.ajhg.2013.04.023 . Accessed 2021-05-29

[61] Geleta, M.: Unsupervised learning with applications in genomics. BS thesis, Universitat Politécnica de Catalunya (2021)

[62] Cover, T.M., Thomas, J.A.: Elements of Information Theory. Wiley series in telecommunications. Wiley, New York (1991)

[63] Nielsen, R., Akey, J.M., Jakobsson, M., Pritchard, J.K., Tishkoff, S., Willerslev, E.: Tracing the peopling of the world through genomics. Nature 541(7637), 302–310 (2017) 10.1038/nature21347. Accessed 2021-05-26

[64] Lan, D., Tobler, R., Souilmi, Y., Llamas, B.: Genozip: a universal extensible genomic data compressor. Bioinformatics (2021) 10.1093/bioinformatics/btab102

[65] Collet, Y., Kucherawy, M.: Zstandard compression and the application/zstd media type. RFC 8478 (2018)

[66] Team, B.D.: Blosc: A high-performance, multithreaded compression library. http://www.blosc.org (2021)

[67] Oord, A., Vinyals, O., Kavukcuoglu, K.: Neural Discrete Representation Learning (2018)

[68] Glorot, X., Bengio, Y.: Understanding the difficulty of training deep feedforward neural networks. In: Teh, Y.W., Titterington, M. (eds.) Proceedings of the Thirteenth International Conference on Artificial Intelligence and Statistics. Proceedings of Machine Learning Research, vol. 9, pp. 249–256. PMLR, Chia Laguna Resort, Sardinia, Italy (2010). https://proceedings.mlr.press/v9/glorot10a.html

[69] Kingma, D.P., Ba, J.: Adam: A Method for Stochastic Optimization (2017)

[70] Srivastava, N., Hinton, G.E., Krizhevsky, A., Sutskever, I., Salakhutdinov, R.: Dropout: a simple way to prevent neural networks from overfitting. Journal of Machine Learning Research 15(1), 1929–1958 (2014)

[71] Suarez-Kurtz, G.: Ethnic differences in drug therapy: a pharmacogenomics perspective. Expert Review of Clinical Pharmacology 1(3), 337–339 (2008) 10.1586/17512433.1.3.337 https://doi.org/10.1586/17512433.1.3.337

